# Phytochemical Nrf2 activator attenuates skeletal muscle mitochondrial dysfunction and impaired proteostasis in a preclinical model of musculoskeletal aging

**DOI:** 10.1101/2021.06.11.448143

**Authors:** Robert V. Musci, Kendra M. Andrie, Maureen A. Walsh, Zackary J. Valenti, Maryam F. Afzali, Taylor Johnson, Thomas E. Kail, Richard Martinez, Tessa Nguyen, Joseph L. Sanford, Meredith D. Murrell, Joe M. McCord, Brooks M. Hybertson, Benjamin F. Miller, Qian Zhang, Martin A. Javors, Kelly S. Santangelo, Karyn L. Hamilton

**Author notes:** Correspondence: Robert V. Musci. Unit for Laboratory Animal Medicine, University of Michigan Medical School, Ann Arbor, MI, USA.

## Abstract

Musculoskeletal dysfunction is an age-related syndrome associated with impaired mitochondrial function and proteostasis. However, few interventions have tested targeting two drivers of musculoskeletal decline. Nuclear factor erythroid 2-related factor 2 (Nrf2) is a transcription factor that stimulates transcription of cytoprotective genes and improves mitochondrial function. We hypothesized daily treatment with a Nrf2 activator in Hartley guinea pigs, a model of age-related musculoskeletal dysfunction, attenuates the progression of skeletal muscle mitochondrial dysfunction and impaired proteostasis, preserving musculoskeletal function. We treated 2-month- and 5-month-old male and female Hartley guinea pigs for 3 and 10 months, respectively, with the phytochemical Nrf2 activator PB125 (Nrf2a). Longitudinal assessments of voluntary mobility were measured using Any-Maze™ open-field enclosure monitoring. Cumulative skeletal muscle protein synthesis rates were measured using deuterium oxide over the final 30 days of treatment. Mitochondrial oxygen consumption in permeabilized soleus muscles was measured using *ex vivo* high resolution respirometry. In both sexes, Nrf2a 1) increased electron transfer system capacity; 2) attenuated the disease/age-related decline in coupled and uncoupled mitochondrial respiration; and 3) attenuated declines in protein synthesis in the myofibrillar, mitochondrial, and cytosolic subfractions of the soleus. These improvements were not associated with statistically significant prolonged maintenance of voluntary mobility in guinea pigs. Collectively, these results demonstrate that treatment with an oral Nrf2 activator contributes to maintenance of skeletal muscle mitochondrial function and proteostasis in a pre-clinical model of musculoskeletal decline. Further investigation is necessary to determine if these improvements are also accompanied by slowed progression of other aspects of musculoskeletal decline.

## Introduction

Targeting the age-related processes that underpin chronic diseases to promote “healthy longevity” or extend the healthspan (Kaeberlein *et al*., 2015) is essential to decrease both healthcare (Atella *et al*., 2019) and financial (Goldman *et al*., 2013) burdens imposed by an increasingly aged population. Moreover, preserving health at later ages would allow for individuals to maintain a greater quality of life. There is a growing list of interventions (Bakula *et al*., 2019) that target the hallmarks (Lopez-Otin *et al*., 2013) and pillars (Kennedy *et al*., 2014) of aging. Thus, evaluating these interventions in translational pre-clinical models represents an essential next-step in developing therapeutics for the human population.

The musculoskeletal system is comprised of bones, joints, cartilage, tendon, and skeletal muscle, all of which are physically and biochemically connected (Bonewald *et al*., 2013; DiGirolamo *et al*., 2013). Age-related decline in musculoskeletal function contributes to the health burden associated with aging (Goates *et al*., 2019). Musculoskeletal dysfunction imparts a loss of mobility and independence (Roux *et al*., 2005) and leads to frailty (Walston *et al*., 2006). It also exacerbates comorbidities including cardiometabolic disease (Baskin *et al*., 2015), cancer (Williams *et al*., 2018), and cognitive decline (Ogawa *et al*., 2018); and increases mortality (García-Hermoso *et al*., 2018). There are no established therapeutics to slow musculoskeletal decline (Yoshimura *et al*., 2017). Accordingly, the NIH identified a critical need (PAR-15-190) to “accelerate the pace of development of novel therapeutics… for preventing and treating key health issues affecting the elderly.”

The lack of effective therapeutics for musculoskeletal disorders is partially attributable to the insidious nature of musculoskeletal decline in humans, as well as the absence of animal models that recapitulate the multifactorial processes that drive musculoskeletal decline. The Hartley guinea pig is an outbred guinea pig that develops primary (also considered spontaneous or idiopathic) osteoarthritis (OA) starting at 4 months of age that closely resembles the onset and disease progression in humans (Jimenez *et al*., 1997). By nine months of age, these guinea pigs have diminished mobility. At 18 months of age, the severity of OA renders the guinea pigs up to 50% less mobile (Santangelo *et al*., 2014). Similar to humans with OA (Kemmler *et al*., 2015; Noehren *et al*., 2018), skeletal muscle fiber size and density decrease and type I fibers increase by 15 months in these guinea pigs (Tonge *et al*., 2013; Musci *et al*., 2020), which in turn worsens the disease and contributes to disability in humans (Lee *et al*., 2016). Thus, the Hartley guinea pig represents a potential model to study musculoskeletal deficiencies associated with osteoarthritis, an age-related chronic disease that affects over 30 million US adults (United States Bone and Joint Initiative, 2020), in a compressed amount of time (i.e. 5 to 15 months of age).

The musculoskeletal system is particularly susceptible to age-related declines in cellular function and increases in damage because it is slow to turnover relative to tissues such as liver (Drake *et al*., 2013). Skeletal muscle is post mitotic and turnover of both bone tissue and cartilage is also slow (Vaananen, 1993; Hall, 2012; Relaix *et al*., 2021). Thus, targeting the hallmarks of aging is likely particularly useful in counteracting age-related musculoskeletal dysfunction. For example, targeting mitochondrial dysfunction likely ameliorates not just impaired ATP production but also other, interconnected hallmarks of aging, such as impaired proteostasis (protein homeostasis) (Musci *et al*., 2018). Impaired mitochondrial function is associated with, and precedes, impairments in proteostasis and decrements in skeletal muscle function (Gaffney *et al*., 2018; Gonzalez-Freire *et al*., 2018). Inversely, improvements in proteostatic mechanisms regulating mitochondrial proteome integrity would improve mitochondrial function (Hamilton & Miller, 2017), which would in turn alleviate the energetic constraints that impair adequate cellular function. This cyclical and interconnected relationship highlights the potential efficacy of targeting one hallmark of aging.

Nuclear factor erythroid 2-related factor 2 (Nrf2) is a transcription factor that regulates hundreds of genes involved in adaptation to stress, including those involved in redox homeostasis, mitochondrial energetics, and proteome maintenance (Gao *et al*., 2020). Nrf2 activation leads to the transcription of genes with the antioxidant response element in the promoter regions, including antioxidant genes such as SOD-1, NQO1, and HO-1 (Kobayashi & Yamamoto, 2006), has anti-inflammatory effects (Ahmed *et al*., 2017), and has a role in regulating mitochondrial biogenesis (Piantadosi *et al*., 2008). Transient Nrf2 activation through phytochemical supplementation (Donovan *et al*., 2012; Reuland *et al*., 2013; Kubo *et al*., 2017; Hybertson *et al*., 2019) is a potential therapeutic intervention that could mitigate age-related chronic diseases (Houghton *et al*., 2016). Transiently activating Nrf2 targets several interconnected drivers of aging including macromolecular damage, disrupted redox homeostasis (Reuland *et al*., 2013; Fang *et al*., 2017), inflammation (Kobayashi *et al*., 2016), and impaired proteostasis (Konopka *et al*., 2017). In the NIH-NIA Interventions Testing Program (ITP), treatment with the phytochemical Nrf2 activator Protandim extended median lifespan of male mice (Strong *et al*., 2016).

Given the positive effects of Nrf2 activator (Nrf2a) treatment, we sought to identify the effects of months-long Nrf2 activator treatment in the Hartley guinea pig. We hypothesized Nrf2a treatment would improve skeletal muscle mitochondrial function and mechanisms of proteostasis and attenuate musculoskeletal declines in both male and female guinea pigs.

## Methods

### Husbandry

All procedures were approved by the Colorado State University Institutional Animal Care and Use Committee and were performed in accordance with the NIH Guide for the Care and Use of Laboratory Animals. Dunkin-Hartley guinea pigs were obtained from Charles River Laboratories (Wilmington, MA, USA) at 1- and 4-months of age (mo) for each treatment regimen such that there were 14 male and female guinea pigs in each age and treatment group (total n = 112) (Figure 1). As mentioned, Hartley guinea pigs begin developing knee OA at 4 mo and have severe OA and skeletal muscle and joint phenotypes consistent with aged human musculoskeletal systems by 15 mo (Jimenez *et al*., 1997; Tonge *et al*., 2013; Santangelo *et al*., 2014; Musci *et al*., 2020). Accordingly, we chose these ages to determine if Nrf2a could prevent the onset (short term treatment from 2 to 5 mo) or mitigate the progression (long term treatment from 5 to 15 mo) of musculoskeletal dysfunction (Jimenez *et al*., 1997; Santangelo *et al*., 2014) and skeletal muscle decline (Musci *et al*., 2020) (Figure 1). It is important to note that because knee OA was progressing as these animals age, we cannot discern the effect of age from disease progression or vice-versa. Thus, for any documented effect of age, we must also acknowledge that the effect could be attributed to disease progression.

**Figure 1.**
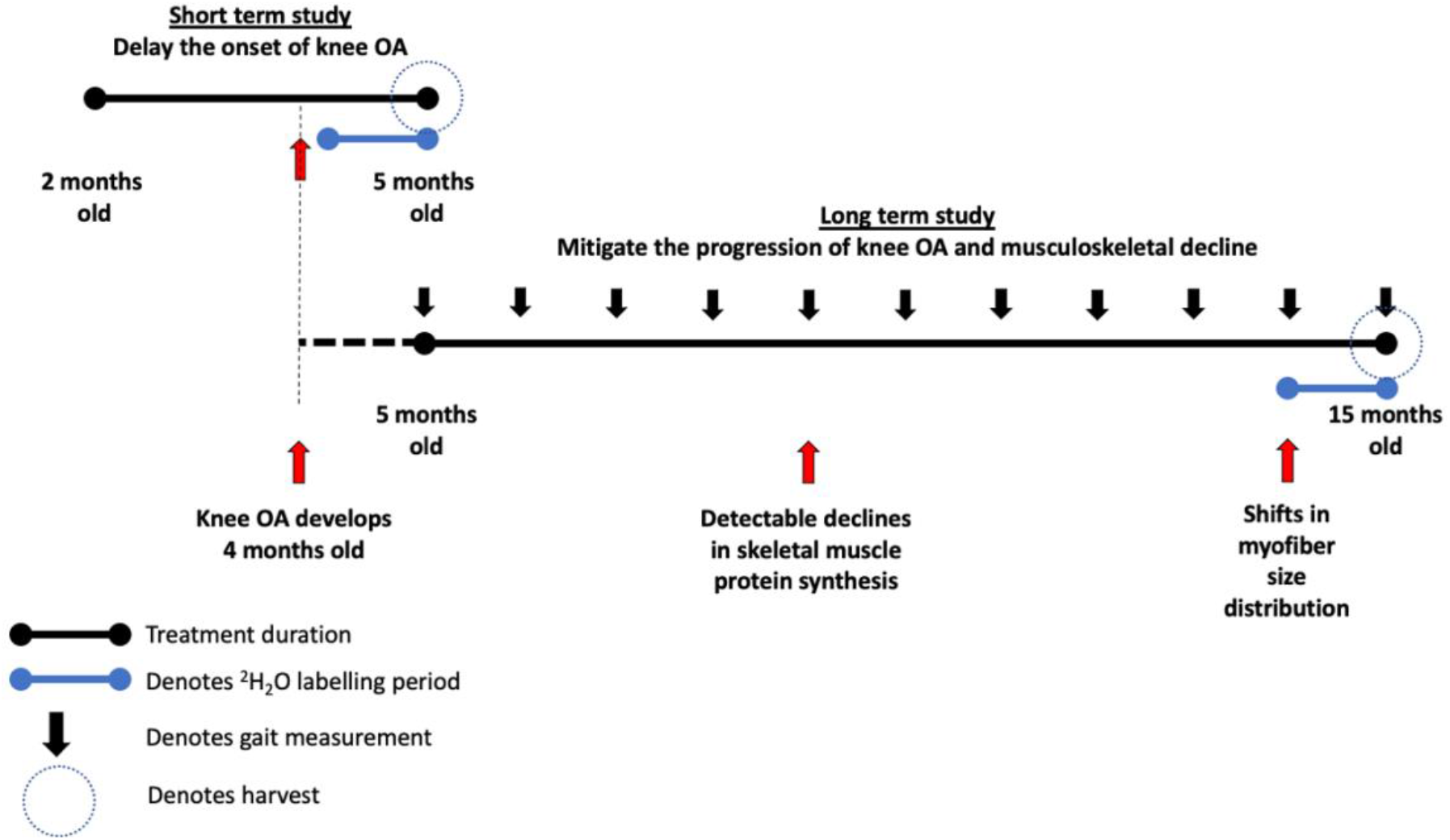
Study Design. There were two cohorts of guinea pigs in this study. The first cohort was treated with Nrf2a or vehicle control from 2 mo to 5 mo, during which knee OA begins developing. The second cohort was treated from 5 mo to 15 mo of age, after which knee OA begins developing and during which detectable declines in musculoskeletal quality arise. In the final 30 days of each study, a bolus I.P. injection of ^2^H_2_O was administered and ^2^H_2_O was mixed in drinking for measurement of protein synthesis. A portion of the soleus was harvested for mitochondrial respirometry assessments. Another portion of the soleus as well as a portion of the gastrocnemius was harvested for isotopic measurements. Comparisons between 5 mo and 15 mo guinea pigs were made between the cohorts at the day of harvest. Longitudinal weight and mobility data were acquired from the second cohort.

Animals were maintained at Colorado State University’s Laboratory Animal Resources housing facilities and were monitored daily by veterinary staff. All guinea pigs were singly-housed in solid bottom cages, maintained on a 12-12 hour light-dark cycle, and provided ad libitum access to food and water. Two control females, two Nrf2a females, one control male, and two Nrf2a males required humane euthanasia prior to final analysis due to underlying issues unrelated to treatment (final n = 105). Gross necropsy findings by veterinarians did not raise significant concern as the cause of death in these cases were consistent with what would be expected in conventionally raised guinea pigs.

### Measurement of PB125 in OraSweet and in Guinea Pig Plasma using High Performance Liquid Chromatography- Mass Spectrometry (HPLC/MS)

PB125 (Pathways Bioscience, Aurora, CO) is a phytochemical compound comprised of rosemary, ashwagandha, and luteolin powders which contain the three active ingredients carnosol (CRN), withaferin A (WFA), and luteolin (LUT) at a mixed ratio of 15:5:2 by mass, respectively (Hybertson *et al*., 2019). Prior to treatment initiation, plasma levels of each activate ingredient were measured 15, 30, 45, 60, 90, and 120 min post dosing at concentrations of 8, 24, and 48 mg/ml (Supplemental Figures 1A – 1C), which corresponds with a dosage of 250, 750, and 1250 PPM. Compound stability in OraSweet was assessed both at room temperature and 4 °C. Reference standards of LUT and WFA were purchased from Sigma Aldrich (St. Louis, MO). CRN was purchased from Cayman Chemical (Ann Arbor, MI). All other reagents were purchased from Thermo Fisher Scientific (Waltham, MA). HPLC grade methanol was used for preparation of all solutions. Samples were analyzed at the Nathan Shock Core Analytical Pharmacology Core at the University of Texas Health Medical School.

The liquid chromatography tandem mass spectrometry (LC/MS/MS) system consisted of a Shimadzu SIL 20A HT autosampler, LC-20AD pumps (2), and an AB Sciex API 4000 tandem mass spectrometer with turbo ion spray. The LC analytical column was an ACE C8 (50 x 3.0 mm, 3 micron) purchased from Mac-Mod Analytical (Chaddsford, PA). Mobile phase A contained 0.1% formic acid dissolved in water. Mobile phase B contained 0.1% formic acid dissolved in 100% HPLC grade acetonitrile. The LC Gradient was: 0 to 2 min, 25% B; 2 to 6 min, linear gradient to 99% B; 6 to 10 min, 99% B; 10 to 10.01 min, 99% to 25% B min; 10.1 to 12 min, 25%B. LUT and CRN were detected in negative mode using these transitions: 285 to 132.9 m/z and 329 to 285 m/z, respectively. WFA was detected in positive mode at the transition of 471 to 281 m/z.

LUT, CRN, and WFA stock solutions were prepared in methanol at a concentration of 1 mg/ml and stored in aliquots at -80 °C. Working stock solutions of each drug were prepared each day from the super stock solutions at a concentration of 100 μg/ml, 10 μg/ml, and 1 μg/ml which were used to spike the calibrators.

Dosages of PB125 in OraSweet were diluted 1000x in 70% ethanol. Calibrator samples were prepared daily by spiking blank OraSweet to achieve final concentrations of 0, 30.4, 152, 760, and 2280 µg/ml. The calibrators were then diluted 1000x in 70% ethanol. The samples were transferred to injection vials and 10 µl was injected into the system. Each drug was quantified by comparing the peak area ratios for each dosage sample against a linear regression of calibrator peak area ratios. The concentration of each drug was reported as µg/ml. Because we prepared weekly allotments of PB125 in Orasweet, we verified the stability of PB125 suspended in OraSweet stored in 4°C for one week (Supplemental Figure 1D).

LUT, CRN, and WFA were also quantified in guinea pig plasma. The transitions used were the same as the OraSweet dilutions. Calibrator samples were prepared daily by spiking blank plasma to achieve final concentrations of 0, 5, 10, 25, 50, 100, 500, 1000, and 5000 ng/ml. Calibrators were left to sit for 5 min after spiking. Briefly, 0.1 mL of calibrator and unknown plasma samples were mixed with 1.0 ml of chilled ethanol, vortexed vigorously, and then centrifuged at 17,000 g for 5 min at 25 °C. The supernatants were transferred to 1.5 ml microcentrifuge tubes and dried to residue under a nitrogen stream. The residues were then redissolved in 60 µL of 50/50 mobile phase A/mobile phase B and were centrifuged 5 min at 17,000 g. The samples were transferred to injection vials and 15 µL was injected into the LC/MS/MS. Each drug was quantified by comparing the peak area ratios for each unknown sample against a linear regression of calibrator peak area ratios. The concentration of LUT, CRN, and WFA were expressed as ng/mL plasma (Supplemental Figure 1A – C).

### Treatment, euthanasia, and tissue acquisition

Based on the analysis conducted at the NSC Analytical Pharmacology Core (Supplemental Figure 1A – C), we selected a dosage of 8 mg/kg of bodyweight, which corresponds to 250 PPM, about 2.5x the dose of PB125 mice in the NIA ITP receive (https://www.nia.nih.gov/research/dab/interventions-testing-program-itp/compounds-testing). This dose was adequate to stimulate Nrf2 activation based on an increase in Nrf2 protein content in the gastrocnemius in a subset of both male and female guinea pigs (Supplemental Figure 1E). *Nrf2* contains an antioxidant response element (ARE) promoter region, which activated Nrf2 proteins bind to upon activation and translocation into the nucleus. Because we were interested in long term effects of Nrf2 treatment (Miller *et al*., 2016), we measured protein concentration instead of mRNA transcript concentration of a downstream Nrf2 target. Additionally, the last dose of the Nrf2 activator was 24 h prior to harvest, which precludes from measuring transcriptional responses to the PB125 treatment. After a one-month acclimation to housing conditions, male and female guinea pigs in each age group (2 or 5 months) were randomized to receive a daily oral dose of 8.0 mg/kg bodyweight of PB125 (Nrf2a) suspended in OraSweet (Perrigo, Dublin, Ireland) or an equivalent volume of OraSweet only (CON). Following established protocol, guinea pigs were given a subcutaneous injection of 0.9% saline enriched with 99% deuterium (^2^H_2_O) equivalent to 3% of their body weight 30 days prior to euthanasia (Musci *et al*., 2020). Drinking water was enriched to 8% ^2^H_2_O for the purpose of maintaining ^2^H_2_O enrichment of the body water pool during the 30-day labelling period. At the time of harvest, the guinea pigs were 5 mo (after 3 months of treatment) or 15 mo (after 10 months of treatment of age). In accordance with the standards of the American Veterinary Medical Association, animals were anesthetized with a mixture of isoflurane and oxygen; thoracic cavities were opened and blood was collected via direct cardiac puncture. Whole blood was centrifuged (1200 *g*, 4 °C, 15 min) to separate plasma, which was frozen at -80 °C until further analysis. After blood collection, the anesthetized animals were transferred a chamber filled with carbon dioxide for euthanasia.

Upon euthanasia, the right leg of the guinea pig was promptly removed for the excision of the soleus muscle. A portion of the right soleus muscle (∼40 mg) was harvested and placed in BIOPS preservation buffer (2.77 mM CaK2-EGTA, 7.23 mM K2-EGTA, 20 mM imidazole, 20 mM taurine, 50 mM K-MES, 0.5 mM dithiothreitol, 6.56 mM MgCl2, 5.77 mM ATP, and 15 mM phosphocreatine, adjusted to pH 7.1) containing 12.5 µM blebbistatin to inhibit muscle contraction (Pesta & Gnaiger, 2011). The rest (∼70 mg) of the soleus was frozen in liquid nitrogen and used for other analyses. After excision of the soleus, at least 70 mg of the right gastrocnemius was collected and frozen immediately in liquid nitrogen. Both soleus and gastrocnemius muscles were trimmed of tendons and connective tissue and weighed. Bone marrow was also harvested in saline from the humeri.

### Mitochondrial respirometry

After the soleus was placed in BIOPS, the muscle fibers were prepared for high resolution respirometry as follows. Mechanical permeabilization occurred on ice using forceps to separate the fibers. After mechanical permeabilization, fibers underwent chemical permeabilization for 30 min in BIOPS with 12.5 µM blebbistatin and 50 µg/mL saponin, followed by a 15 min rinse in BIOPS. Approximately 2.0 mg (wet weight) of muscle fibers were placed in mitochondrial respiration medium (MiR05, 0.5 mM EGTA, 3 mM MgCl_2_6H_2_O, 20 mM Taurine, 15 mM Na_2_Phosphocreatine, 20 mM Imidazole, 0.5 mM Dithiothreitol, and 50 mM K^+^ -MES at pH 7.1) in an Oxygraph-2k (O2K) (Oroboros, Innsbruck, Austria) for high resolution respirometry. To control for oxygen flux at higher concentrations of oxygen, each morning of respirometry analysis, we conducted high oxygen concentration calibrations at 450, 350, 250, and 167 (i.e., concentration of room air) nmol/ml O_2_ (Pesta & Gnaiger, 2011). During the experiments, oxygen concentrations were maintained between 225 – 450 nmol/ml O_2_. High resolution respirometry measurements were performed in duplicate using two different protocols. Please refer to Supplemental Table 1 for a detailed explanation of the protocols.

The first protocol (SUIT 1) was an ADP titration protocol to determine ADP sensitivity (Km) and maximal oxidative capacity (Vmax) under Complex I supported respiration. We measured Complex I supported leak respiration (State 2_[PGM]_) with the addition of 10 mM glutamate, 0.5 mM malate, and 5 mM pyruvate. Upon acquisition of State 2_[PGM]_, we titrated progressively greater concentrations of ADP from 0.1 mM, 0.175 mM, 0.25 mM, 1 mM, 2 mM, 4 mM, 8 mM, 12 mM, 20 mM, to 24 mM (State 3_[PGM]_), awaiting steady-state oxygen flux prior to adding the subsequent titration to determine Complex I linked ADP Vmax and apparent Km (i.e. ADP sensitivity). After the ADP titration was completed, we added 5 mM cytochrome C to test mitochondrial membrane integrity. After cytochrome C addition, we added 10 mM succinate to acquire maximal Complex I and II supported coupled respiration (State 3_[PGM + S]_). We then added 0.5 µM FCCP sequentially until there was no increase in respiration to determine the capacity of the electron transport system to consume oxygen, or maximal uncoupled respiration (ETS_[CI-CIV]_). Finally, we added 5 µM rotenone to measure maximal uncoupled respiration with the inhibition of Complex I (ETS_[CII-CIV]_), followed by 2.5 µM Antimycin A to measure residual oxygen consumption (ROX).

The second protocol (SUIT 2) measured oxygen consumption while simultaneously measuring ROS production by using the fluorometer attachment of the O2K (Robinson *et al*., 2019) and addition of 10 µM Amplex Red, 1 U/ml horseradish peroxidase, and 5 U/ml superoxide dismutase. We then measured fatty acid supported leak respiration by adding 10 mM glutamate, 0.5 mM malate, 5 mM pyruvate, and 0.2 mM octanoylcarnitine (State 2_[PGM + Oct]_) and 10 mM succinate (State 2_[PGM + Oct + S]_). After stimulating maximal leak respiration, we added submaximal boluses of ADP (0.5 mM: (State 3_[Sub + 0.5D]_) and 1 mM: State 3_[Sub + 1.0D]_), followed by a saturating bolus of ADP (6.0 mM: State 3_[Sub + 6.0D]_). We added 5 mM cytochrome C to test mitochondrial membrane integrity. We set a cytochrome C control factor threshold of 0.25. We set this threshold based on the presence of a negative linear relationship between the cytochrome C control factor and State 3 respiration in the SUIT 2 protocol. Upon eliminating respirometry trials that had a cytochrome C control factor of greater than 0.25, the negative linear relationship no longer existed and all samples included in analysis were not biased by over-permeabilization, which is what the cytochrome C control factor approximates (Pesta & Gnaiger, 2011) (Supplemental Figure 4 A - C). We then added 5 µM rotenone to determine maximal coupled respiration in the absence of Complex I (State 3_[Sub + D - CI]_) followed by sequential titrations of 0.5 µM FCCP until respiration no longer increased to determine maximal fatty acid supported uncoupled respiration (ETS_[Sub + D – CI]_). and added 2.5 µM antimycin A to measure ROX. The respiratory control ratio (RCR: State 3/State 2), which is an index of mitochondrial efficiency was also evaluated.

### Protein isolation and fractionation

The gastrocnemius and soleus muscles were homogenized and fractionated following established laboratory protocols (Drake *et al*., 2013; Miller *et al*., 2013; Groennebaek *et al*., 2018; Sieljacks *et al*., 2019; Musci *et al*., 2020). Briefly, tissues (20 – 50 mg) were homogenized at 1:10 in isolation buffer (100 mM KCl, 40 mM Tris HCl, 10 mM Tris Base, 5 mM MgCl2, 1 mM EDTA, 1 mM ATP, pH – 7.5) with phosphatase and protease inhibitors (HALT Thermo Scientific, Rockford, IL, USA) using a tissue homogenizer (Bullet Blender, Next Advance Inc., Averill Park, NY, USA) with zirconium beads (Next Advance Inc., Averill Park, NY, USA). After homogenization, subcellular fractions were isolated via differential centrifugation as previously described (Musci *et al*., 2020). Once fractionated pellets were isolated and purified, 250 µl 1 M NaOH was added and pellets were incubated for 15 min at 50 °C and 900 RPM.

### DNA extraction

Approximately 100 ng/µL of total DNA was extracted from 20 mg tissue (QiAMP DNA mini kit Qiagen, Valencia, CA, USA). DNA from bone marrow was extracted from the bone marrow suspension and centrifuged for 10 min at 2000 *g*, yielding approximately 100 ng/µL. *Sample preparation and analysis via GC/MS: Proteins*

Protein subfractions were hydrolyzed in 6 M HCl for 24 hours at 120 °C after which the hydrolysates were ion-exchanged, dried *in vacuo*, and then resuspended in 1 mL of molecular biology grade H_2_O. Half of the suspension was derivatized with 500 µL acetonitrile, 50 µL 1 M K_2_HPO_4_, and 20 µl of pentafluorobenzyl bromide and incubated at 100 °C for 60 min. Derivatives were extracted into ethyl acetate and the organic layer was transferred into vials which were then dried under nitrogen. Samples were reconstituted in ethyl acetate (200 µL – 700 µL).

The derivative of alanine was analyzed on an Agilent 7890A GC coupled to an Agilent 5977A MS as previously described (Robinson *et al*., 2011; Drake *et al*., 2013; Miller *et al*., 2013; Groennebaek *et al*., 2018; Miller *et al*., 2019; Sieljacks *et al*., 2019; Musci *et al*., 2020). The newly synthesized fraction (f) of proteins was calculated from the true precursor enrichment (p) based upon plasma analyzed for ^2^H_2_O enrichment and adjusted using mass isotopomer distribution analysis (Busch *et al*., 2005). Protein synthesis of each subfraction was calculated as the fraction of deuterium-labeled over unlabeled alanine proteins over the entire labeling period (30 days) and expressed as the fractional synthesis rate (FSR). Thus, we divided fraction new by our labeling period (30 days) and multiplied by 100 to express FSR as %/day. Our isotope approach and analysis followed the established procedures detailed in this Core of Reproducibility in Physiology publication (Miller *et al*., 2020).

### Sample preparation and analysis via Gas Chromatography/Mass Spectroscopy (GC/MS): Body water

80 µL of plasma was placed into the inner well of an o-ring cap that was screwed to tube and inverted on a heating block overnight at 100 °C. After incubation, 2 µL of 10 M NaOH and 20 µL of acetone were added to the samples and ^2^H_2_O standards (0 – 20%) and capped immediately, vortexed, and incubated at room temperature overnight. Samples were extracted with 200 µL hexane and the organic layer was transferred through pipette tips filled with anhydrous Na_2_SO_4_ into GC vials and analyzed via EI mode using a DB-17MS column.

### Sample preparation and analysis via GC/MS: DNA

Incorporation of ^2^H into purine deoxyribose (dR) of DNA was measured follow procedures already described (Busch *et al*., 2007; Miller *et al*., 2012; Drake *et al*., 2013; Drake *et al*., 2014). DNA that was isolated from tissue and bone marrow were hydrolyzed with nuclease S1 and potato acid phosphatase at 37 °C shaking at 150 RPM overnight. These hydrolysates were derivatized with pentafluorobenzyl hydroxylamine and acetic acid and incubated at 100 °C for 30 min. After incubation, samples were acetylated with acetic anhydride and 1-methylimidazole. Dichloromethane was added, mixed and then extracted, dried *in vacuo*, and resuspended in ethyl acetate, and analyzed by GC/MS as previously described (Busch *et al*., 2007; Miller *et al*., 2012; Drake *et al*., 2014; Drake *et al*., 2015). The fraction new was calculated by dividing deuterated dR of the muscle tissue by the bone marrow of the same animal, which represents a fully turned-over cell population, and thus indicative of precursor enrichment (Miller *et al*., 2012; Miller *et al*., 2014; Drake *et al*., 2015).

### Assessing protein synthesis related to mechanisms of proteostasis

To evaluate protein synthesis related to protein maintenance versus new cell proliferation (new DNA), we calculated the ratio of protein synthesis to DNA synthesis (Drake *et al*., 2014; Miller *et al*., 2014; Drake *et al*., 2015; Hamilton & Miller, 2017). Increases in PRO:DNA is indicative of a greater proportion of protein synthesis related to protein turnover to maintain the proteome, with less dedicated to proliferation.

### Protein content

Western blotting was used to measure relative content of Nrf2 and OXPHOS proteins in a subset of tissues. 50-70 mg portions of gastrocnemius and 30 mg portions of soleus (n=9 per treatment group) were powdered under liquid nitrogen and homogenized in a Bullet Blender with zirconium beads and 1.0 mL of radioimmunoprecipitation assay (RIPA) buffer (150 mM NaCl, 0.1 mM EDTA, 50 mM Tris, 0.1% sodium deoxycholate, 0.1% SDS, 1% Triton X-100, pH = 7.50) with HALT protease inhibitors. Samples were reduced (50 µL of B-mercaptoethanol) and heated at 50 °C for 10 min. Approximately 10 µg of protein was loaded into a 4% - 20% Criterion pre-cast gel (Bio-Rad, Hercules, CA, USA) and resolved at 120 V for 120 min. The proteins were then transferred to a PVDF membrane at 100 V for 75 min in transfer buffer (20% w/v methanol, 0.02% w/v SDS, 25 mM Tris Base, 192 mM glycine, pH 8.3). Protein transfer to membrane was confirmed with ponceau stain. Membranes were then blocked and incubated with primary antibodies against Nrf2 (Santa Cruz 13032) and total OXPHOS proteins (Abcam 110413) diluted to 1:500 on a shaker overnight in 4 °C. Membranes were rinsed and then incubated with appropriate secondary antibodies (Santa Cruz 2004 and 2005, respectively) diluted to 1:10,000 for 45 min at room temperature. Protein carbonyls were measured by following the protocol in the commercially available OxiSelect Protein Carbonyl Immunoblot Kit (Cell Biolabs STA-308) as previously performed (Konopka *et al*., 2015; Konopka *et al*., 2017). After the membranes were rinsed, SuperSignal West Dura Extended Duration Substrate (Thermo Fisher 34075) was applied and the membranes were subsequently imaged using a FluorChem E Chemiluminescence Imager (Protein Simple, San Diego, CA, USA). Analysis of densitometry was completed using AlphaView SA Software. Units are expressed as density of primary antibody relative to density of ponceau staining.

### Mobility

Animals were acclimated over a 2-week period, before the onset of the study, to an open circular field behavior monitoring system (ANY-maze™, Wood Dale, IL) to assess voluntary physical mobility. Animals’ activities were recorded, and data were collected for 10 consecutive minutes on a monthly basis throughout the study. Videos were analyzed for the following parameters: total distance traveled (m), average speed (m/s), time mobile (s), % time mobile, time in hut (s), % time in hut, and average moving speed (m/s). Because musculoskeletal degradation causes immobility in Hartley guinea pigs, Kaplan Meier curves combined with log rank and Gehan-Breslow-Wilcoxon tests were utilized to assess the probability of sustained voluntary mobility throughout the 10-month study period of the “long term” study. The “event” was task noncompliance and defined as number of weeks into the study until an animal did not move (i.e., zero distanced traveled when exposed to the open circular field). Remaining individuals that maintained mobility throughout the entire 40-week study duration were censored at the 40-week study endpoint.

### Statistics

For mitochondrial respirometry, in line with best practices, technical replicates were averaged. The average variability between these technical replicates in this study was 18%, which is standard according to the literature (Jacques *et al*., 2020). Apparent Km and Vmax values were determined using Michaelis-Menten kinetics in Prism 9.0 (La Jolla, California, USA). For evaluating growth rates, a non-linear Gompertz growth line was fit to the change in body mass over time (i.e. the rate of growth). The rate of growth, k, was compared between treatment and control within in each sex. For respirometry, isotopic measures, and Western blots, three-way ANOVAs were used to measure the main effects of sex, disease/age, and Nrf2a treatment. Post-hoc analyses were performed using Bonferroni’s post-hoc test.

To determine the effect of Nrf2a on disease/age-related changes in mitochondrial respiration and protein synthesis when a significant effect of disease/age was detected, we conducted a subset analysis using a one-way ANOVA with a Dunnett’s post-hoc test comparing 15 mo treated and untreated guinea pigs to 5 mo untreated guinea pigs.

To assess treatment and sex effects in PRO:DNA of 15 mo guinea pigs, we used a Two-Way ANOVA. We excluded 5 mo guinea pigs from this analysis because of the significantly greater DNA fraction new in 5 mo guinea pigs compared to 15 mo guinea pigs, which makes the age-related comparison of PRO:DNA less relevant. Due to sample loss during processing of bone marrow, there is a reduced sample size for DNA fraction new outcomes, which also affected PRO:DNA.

For Western blots, a subset (n=9) of guinea pigs were randomly selected for analysis. Because this study was a secondary project within a larger study with a different primary outcome, we did not design this study to be powered to detect differences in mitochondrial respiration and protein synthesis at a p-value <0.05. While we set statistical significance *a priori* at p<0.05, we also report differences with p<0.10 to highlight potential directions for future studies. Data are presented as mean +/-SD. All statistics were performed in Prism 9.0 (La Jolla, California, USA).

## Results

Growth rate (k-curves) of the group treated with PB125 (Nrf2a) did not significantly differ from the control group (CON) as measured by changes of body weight throughout treatment (p>0.70 for both sexes) and body mass at harvest (p>0.70) (Supplemental Figures 2A - C). Skeletal muscle DNA synthesis, which is reflective of proliferation of various cell types within the skeletal muscle niche, also was not different between Nrf2a and CON (Supplemental Figure 2D - E). Moreover, there were no differences in absolute or relative skeletal muscle mass between Nrf2a and CON (Supplemental Figure 3).

### Disease/age-related declines in mitochondrial respiration are not sex-specific in Hartley guinea pigs

Because mitochondrial respiratory capacity has never been measured in permeabilized skeletal muscle fibers from Hartley guinea pigs, we first evaluated disease/age- and sex-differences in mitochondrial respiration. As a reference, refer to Supplemental Table 1 for a glossary and detailed titration data for respiratory state mentioned below. Eleven of 210 trials were excluded due to over-permeabilization (cytochrome C control factor > 0.25; Supplemental Figures 4A - B). Maximal coupled (State 3_[CI-CIV]_) (Figure 2A) and uncoupled (Electron Transport System (ETS) _[CI-CIV]_) (Figure 2B) respiration were significantly greater in males (p=0.006; p=0.002, respectively). Uncoupled Complex II-IV (ETS_[CII-CIV]_) supported respiration was greater in males than females (Figure 2C) (p=0.002). However, there was no difference in mitochondrial efficiency (RCR) between sexes (Figure 2D). Males have greater fatty acid supported coupled respiration at sub-saturating (1 mM ADP) and saturating (6 mM ADP) concentrations of ADP (p=0.024 and p=0.018, respectively) (Figures 3B - C).

**Figure 2.**
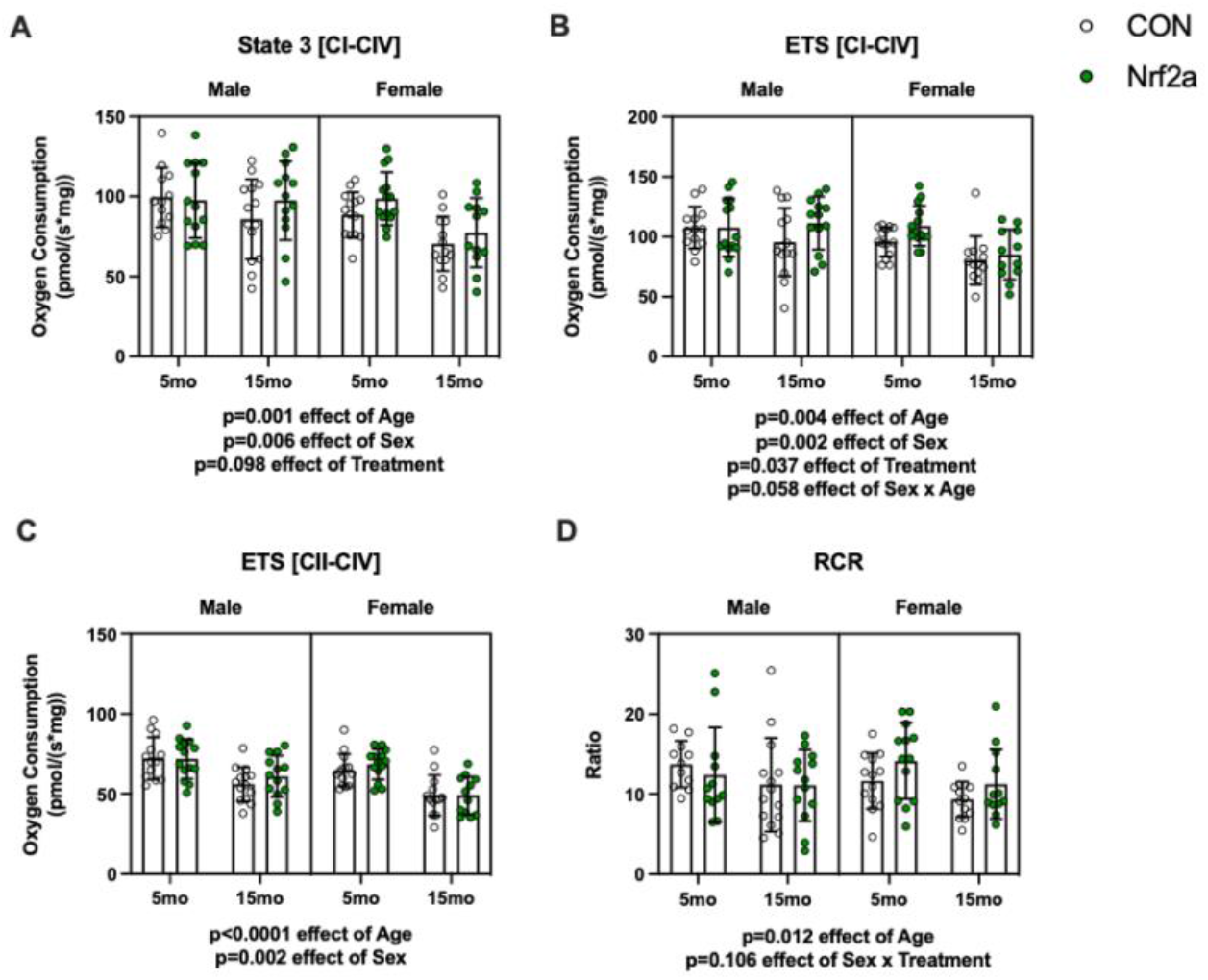
Age-, Sex-, and Treatment-related differences in mitochondrial respiration. There was a significant negative effect of Age on State 3_[CI-CIV]_ respiration (p=0.001). Female guinea pigs had lower levels of respiration compared to males (p=0.006). Treatment did not significantly increase respiration (p=0.098) **(A)**. Electron transport system capacity (ETS_[CI-CIV]_) significantly decreased with age (p=0.004) and was lower in females (p=0.002). Nrf2a treatment increased respiration (p=0.037). The interaction effect between Sex and Age was insignificant (p=0.058) **(B)**. There was a significant decrease in Complex II – IV uncoupled respiration with age (p<0.0001), but there was no effect of Treatment. Female guinea pigs had lower respiration compared to male guinea pigs (p=0.002 effect of Sex) **(C)**. Mitochondrial efficiency (RCR) decreased with age (p=0.012) **(D)**.

Disease/age had a negative effect on several aspects of mitochondrial function in both male and female guinea pigs. 15 mo male and female guinea pigs had lower coupled (State 3_[PGM+S]_) (Figure 2A) and uncoupled (ETS_[CI-CIV]_) (Figure 2B) respiration (p=0.001, p=0.004, respectively). There was also a disease/age-related decline (p<0.0001) in uncoupled respiration without Complex I support (ETS_[CII-CIV]_) (Figure 2C). Disease/age had no effect on fatty acid oxidation supported respiration at sub-saturating levels ADP (Figures 3A – B), though 15 mo guinea pigs had lower fatty acid oxidation supported respiration at saturating levels of ADP (Figure 3C; p=0.056). Mitochondrial efficiency (RCR) also decreased as a result of disease/age (p=0.012) (Figure 2D).

**Figure 3.**
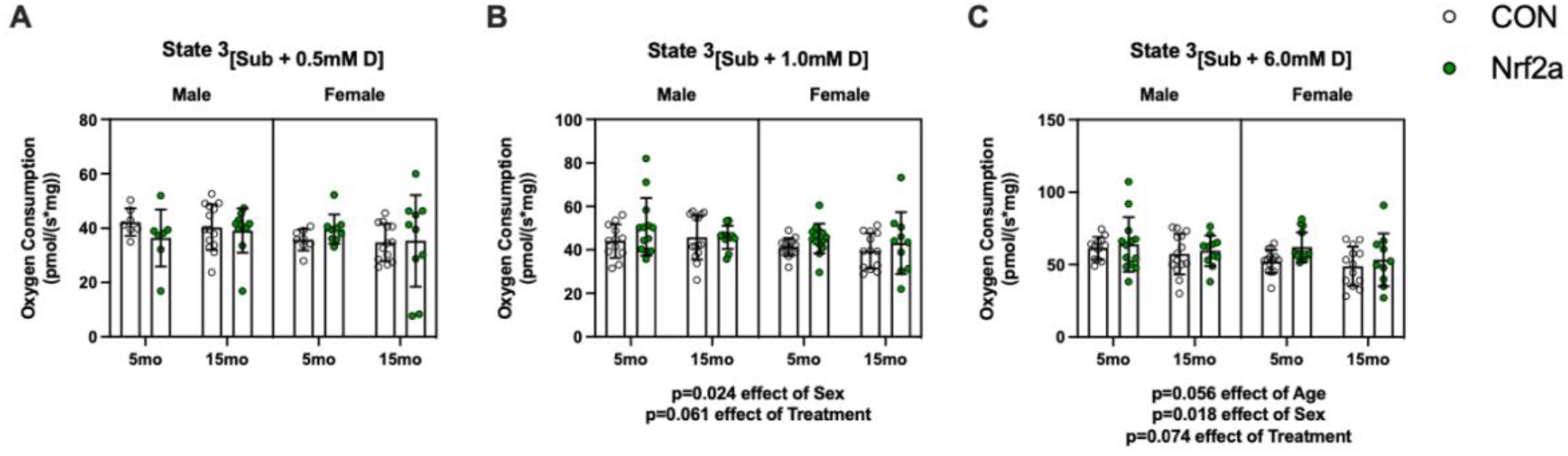
Fatty acid supported respiration. There was no difference in fatty acid supported respiration with 0.5 mM ADP between Sex, Age, or Treatment groups **(A)**. Fatty acid supported respiration with 1.0 mM was lower in females (p=0.029 effect of Sex), but the effect of Treatment was insignificant (p=0.086) **(B)**. At saturating amounts of ADP (6.0 mM), female guinea pigs had lower fatty acid supported respiration compared (p=0.030 effect of Sex), though there was not a significant difference between 5 mo and 15 mo guinea pigs (p=0.058 effect of Age). Nrf2a treatment did not have a significant effect on respiration (p=0.098) **(C)**.

### Nrf2a improves mitochondrial respiration in both male and females

Nrf2a improved several components of mitochondrial respiration in both 5 mo and 15 mo guinea pigs, and in both males and females. Nrf2a did not significantly enhance coupled respiration (State 3[PGM+S]) in male and female guinea pigs (p=0.098) (Figure 2A), but did significantly increase electron transport system (ETS) capacity (ETS_[CI-CIV]_) (Figure 2B; p=0.037). However, Nrf2a did not influence uncoupled respiration with Complex I inhibited (ETS[CII-CIV]) (Figure 2C).

Nrf2a did not significantly improve fatty acid supported respiration at sub-saturating (1 mM ADP) and saturating (6 mM ADP) concentrations of ADP (Figures 3B - C; p=0.061, p=0.074, respectively). There was no main effect of Nrf2a on RCR (Figure 2D), a metric of mitochondrial efficiency, or on ROS emission (Supplemental Figure 5).

### Nrf2a has sex specific effects on mitochondrial ADP kinetics

No O2K data from the ADP titration protocol were excluded based on cytochrome C control factors as all values were below 0.25 (Supplemental Figure 4C). We determined ADP kinetics by titrating progressively higher concentrations of ADP with saturating amounts of pyruvate, glutamate, and malate (titration curves found in Supplemental Figure 6E – H). ADP Vmax was greater in both 15 mo male and female guinea pigs compared to 5 mo counterparts (p=0.049) (Figure 4A). In females, ADP Vmax was lower compared to males (p=0.001) (Figure 4A). Guinea pigs that received treatment with the Nrf2 activator (Nrf2a) had a greater Complex I supported ADP Vmax. Post-hoc comparisons indicate that Nrf2a improved ADP Vmax in 5 mo female guinea pigs (p=0.045).

**Figure 4.**
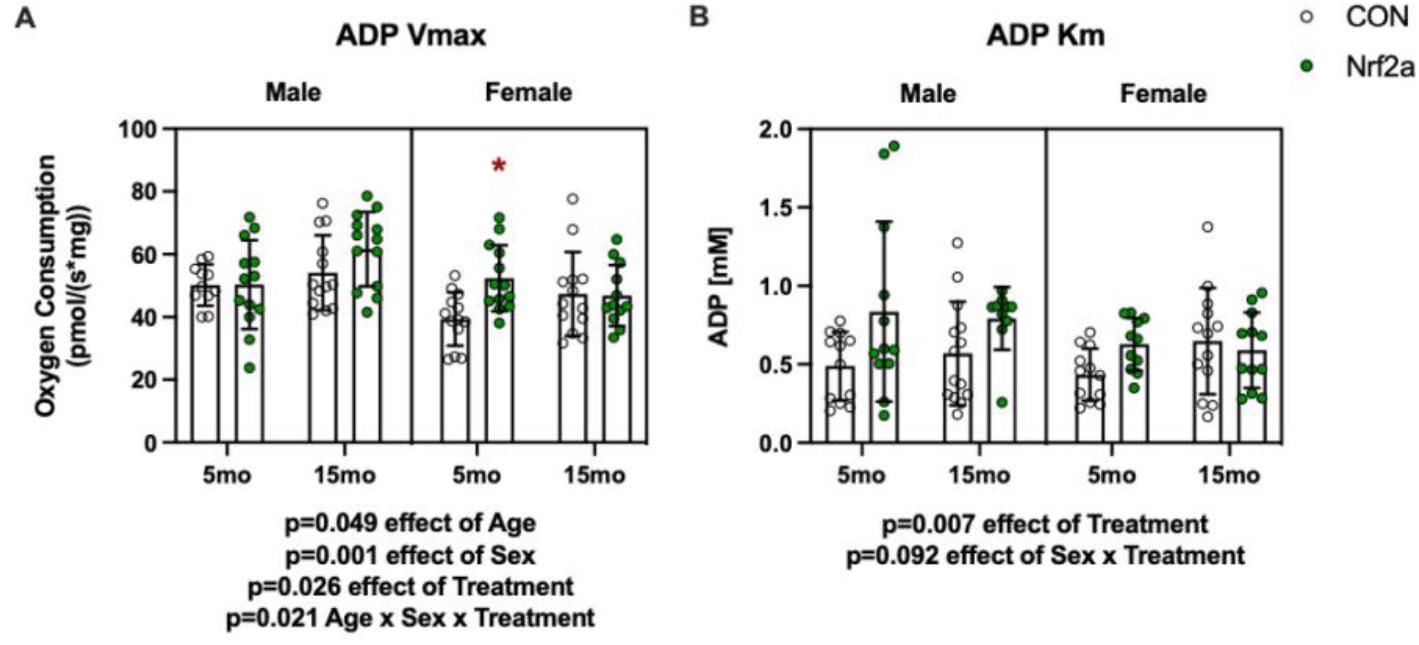
Nrf2a treatment improves ADP Vmax. There was an age-related increase in ADP Vmax (p=0.049), though female guinea pigs had a lower Vmax compared males (p=0.001). Nrf2a significant increased ADP Vmax (p=0.026). Post-hoc analysis revealed Nrf2a 5 mo female had greater ADP Vmax compared to CON 5 mo female guinea pigs (p=0.045) **(A)**. There was a significant increase in ADP Km from Nrf2a treatment (p=0.007). There was an insignificant interaction between Sex and Treatment (p=0.092) **(B)**.

Despite ADP Vmax being greater in 15 mo guinea pigs, there was no effect of disease/age on the apparent Km of ADP (Figure 4B). There were also no differences in Km between sexes. However, Nrf2a did significantly increase the apparent Km (p=0.007) indicating lower ADP sensitivity, though this is likely a consequence of increased ADP Vmax in the absence of changes in respiration rates in sub-saturating amounts of ADP (Supplemental Figure 6). There was non-significant interaction between sex and Nrf2a treatment (p=0.092), indicating that the Nrf2a-mediated decrease in Km may have occurred only in males.

### Nrf2a attenuates age-related declines in mitochondrial respiration

For any main effects of disease/age on mitochondrial respiration, we evaluated if Nrf2a attenuated the age-related changes. That is, where we identified significant differences between 5 mo CON and 15 mo CON guinea pigs, but no differences between 5 mo CON and 15 mo Nrf2a animals, we reported those findings as an attenuating effect of Nrf2a treatment on age-related changes in mitochondrial function. While there was a main positive effect of age on ADP Vmax in the three-way ANOVA (Figure 4A), there was no difference in ADP Vmax between 5 mo and 15 mo CON guinea pigs (p=0.109) in the subsequent one-way ANOVA analysis (Figure 5A). Treated 15 mo guinea pigs, however, had a significantly higher ADP Vmax compared to 5 mo animals (p=0.007) (Figure 4A). Interestingly, this effect was only observed in males (p=0.021) (Figure 5B). While ADP Vmax was greater in 15 mo guinea pigs, 15 mo guinea pigs had a significantly (p=0.024) lower maximal coupled respiration (State 3_[CI-CIV]_) compared to 5 mo counterparts (Figure 5C). Nrf2a, however, prevented that disease/age-related decline (Figure 5C). Maximal uncoupled respiration (ETS_[CI-CIV]_) was also lower between 5 mo and 15 mo CON (Figure 5E), but Nrf2a prevented the decline. Further interrogation revealed that 15 mo females had lower ETS_[CI-CIV]_ compared to their 5 mo counterparts, which Nrf2a attenuated (Figure 5F). Interestingly, when Complex I was inhibited, Nrf2a had no effect on uncoupled respiration (ETS_[CII-CIV]_) and had no effect on the disease/age-related decline in CII-CIV capacity in either 15 mo males (p=0.035) or females (p=0.003) (Figures 5G - H). While the RCR of 15 mo CON guinea pigs were lower compared to 5 mo guinea pigs, RCR was not different between 15 mo Nrf2a treated guinea pigs and 5 mo CON (Figure 5I). However, this occurred only in males where there was a significant difference (p=0.036) between 5 mo and 15 mo CON animals (Figure 5J) but no difference (p=0.151) between 15 mo males treated with Nrf2a compared to 5 mo CON (Figure 5J). There was no difference in RCR between 5 mo and 15 mo females (Figure 5J). Altogether, these data support that Nrf2a can attenuate age related declines in mitochondrial respiration. Interestingly, Nrf2a had no effect on mitochondrial content as assessed by Western blot (Supplemental Figure 7), suggesting that the improvements in mitochondrial function are independent of mitochondrial content in skeletal muscle and may reflect improved mitochondrial quality.

**Figure 5.**
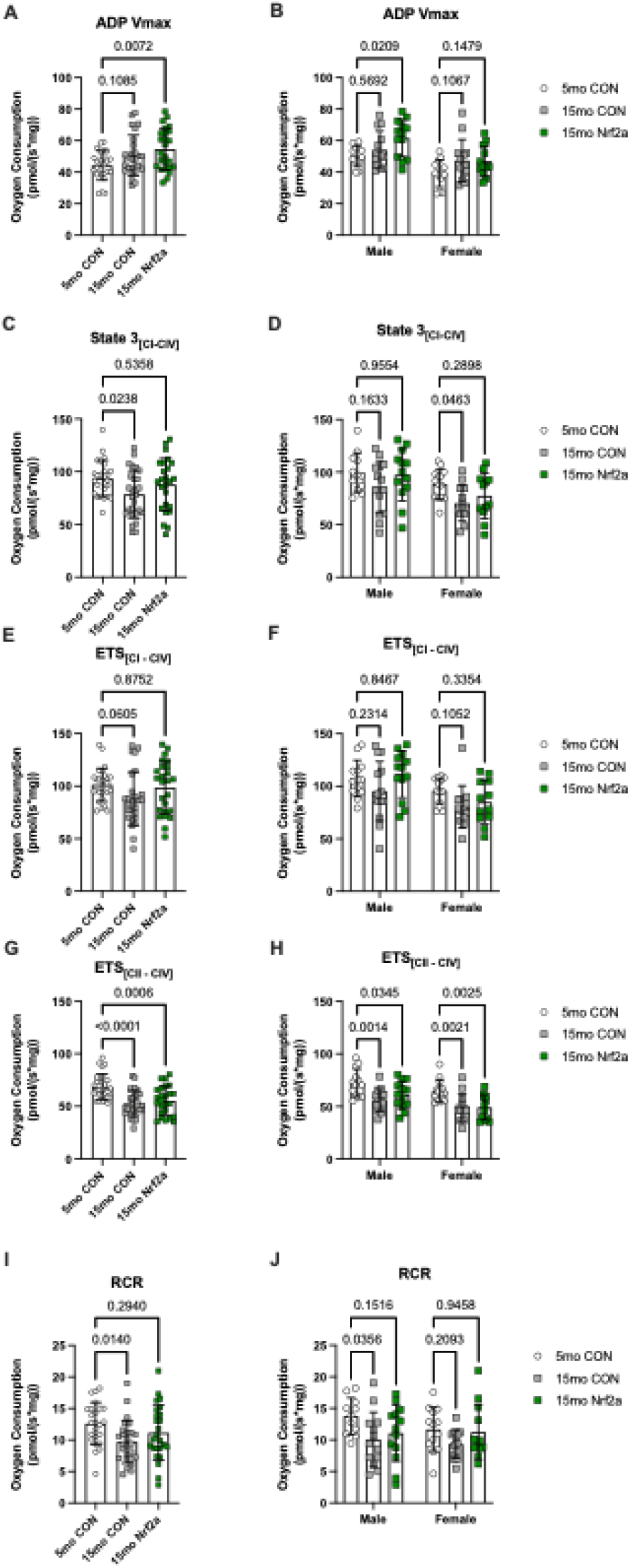
Nrf2a attenuates age-related declines in mitochondrial respiration. There was no difference in ADP Vmax between 5mo and 15mo CON guinea pigs (p=0.109), whereas 15mo Nrf2a guinea pigs had a higher ADP Vmax than 5mo CON guinea pigs (p=0.072) **(A)**. Comparing sex-specific effects, Nrf2a only had a positive effect in male guinea pigs only (p=0.021) **(B)**. There was an age-related decrease (p=0.024) in State 3_[PGM + S]_ between CON guinea pigs, though this difference was attenuated in 15mo Nrf2a guinea pigs (p=0.536) **(C)**. The age-related decline though, was only observed in female guinea pigs (p=0.046), which was attenuated by Nrf2a (p=0.290) **(D)**. Uncoupled respiration ETS_[CI – CIV]_ non-significantly (p=0.061) decreased with age, though Nrf2a attenuated this difference (p=0.875) **(E)**, though there were no significant differences when sex was considered **(F)**. There was a significant decrease (p<0.0001) in ETS_[CII – CIV]_ between 5 mo and 15 mo CON guinea pigs that Nrf2a did not attenuate (p=0.0006) **(G)** in either sex **(H)**.

### Age-sex- and treatment-related effects on skeletal muscle protein synthesis

To determine whether or not Nrf2a-mediated improvement in mitochondrial respiration was linked to improvements in components of proteostasis, we used ^2^H_2_O to measure cumulative protein and DNA synthesis rates over 30 days. There were no differences in fractional synthesis rate (FSR) in either the gastrocnemius or soleus between male and female guinea pigs (Figure 6). There was a disease/age-related decline in the rates of protein synthesis in all subfractions in the soleus and gastrocnemius of both male and female guinea pigs (p<0.01 for all subfractions) (Figure 6). Nrf2a did not have a main effect on FSR in any of the subfractions of either muscle from 5 mo or 15 mo, male or female guinea pigs (Figure 6). However, there was a non-significant interaction between age and Nrf2a (p=0.086) in the myofibrillar subfraction of the soleus of both male and female guinea pigs, suggesting that Nrf2a may have had a positive effect on myofibrillar FSR at 15 mo (Figure 6A).

**Figure 6.**
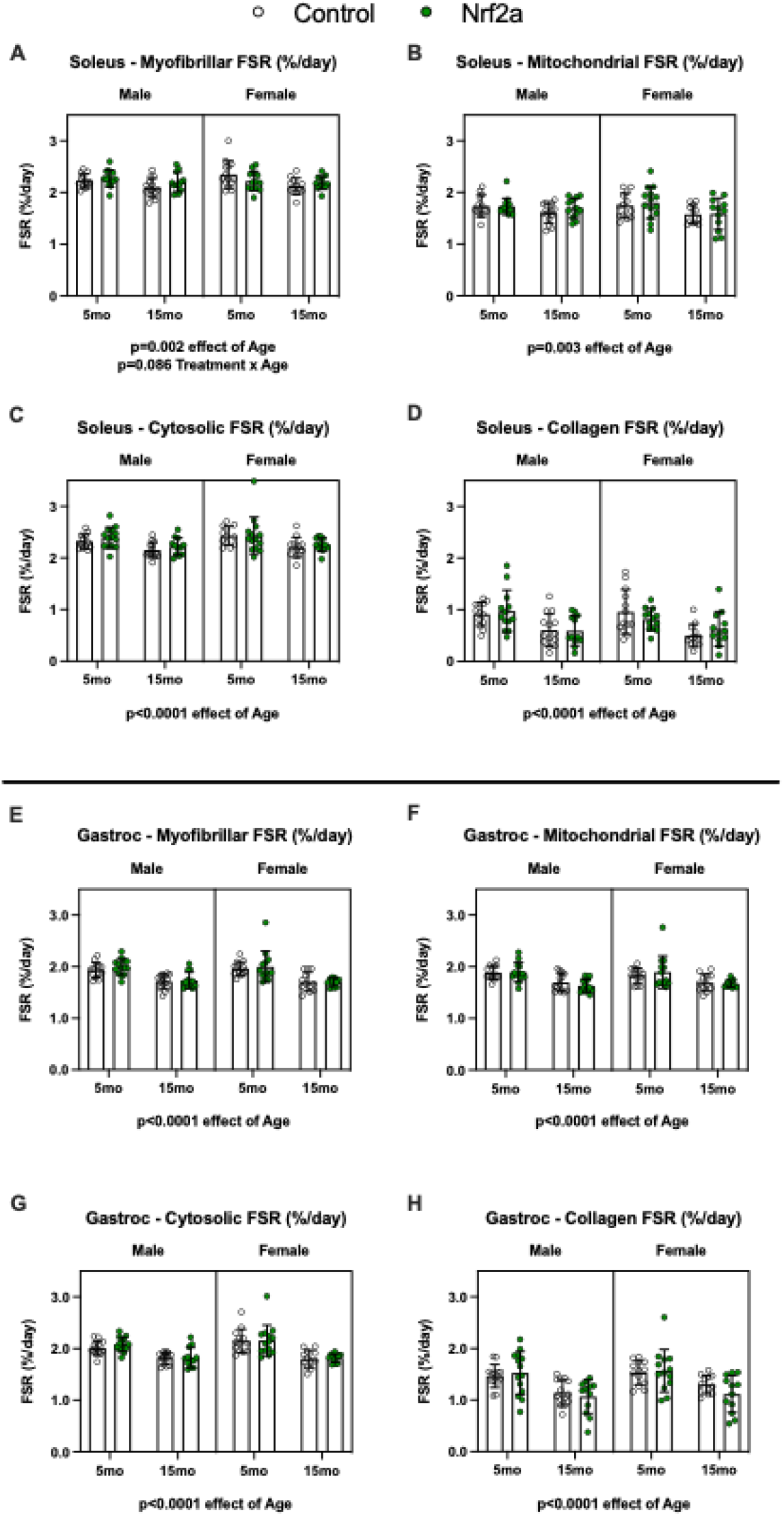
Fractional synthesis rates (FSR) of both the soleus and gastrocnemius subfractions decrease with age. FSR significantly decreased with age in all subfractions of the soleus (p=0.002, p=0.003, p<0.0001, p<0.0001 for myofibrillar **(A)**, mitochondrial **(B)**, cytosolic **(C)**, and collagen **(D)** subfractions, respectively). 15 mo guinea pigs also had a significant decrease in FSR in every subfraction of gastrocnemius as well (p<0.0001 for all subfractions) **(E – H)**.

### Nrf2a mitigates age-related declines protein synthesis

Because there was a disease/age-related decline in protein synthesis rates in all subfractions of both the soleus and gastrocnemius, we sought to determine if Nrf2a prevented any of those declines. Nrf2a attenuated the disease/age-related decline in myofibrillar FSR of the soleus in both males and females (Figures 7A - B). Additionally, Nrf2a attenuated the decline in mitochondrial FSR in the soleus (Figure 7C), but these significant differences were no longer detectable when evaluated in males and females separately (Figure 7D). In the soleus, Nrf2a also mitigated the decline in cytosolic FSR in males only (Figure 7F), but had no effect on the decline in collagen FSR in either sex (Figures 7G - H). In contrast, Nrf2a had no attenuating effect on the disease/age-related decline in protein synthesis in any subfraction of the gastrocnemius (Figure 8).

**Figure 7.**
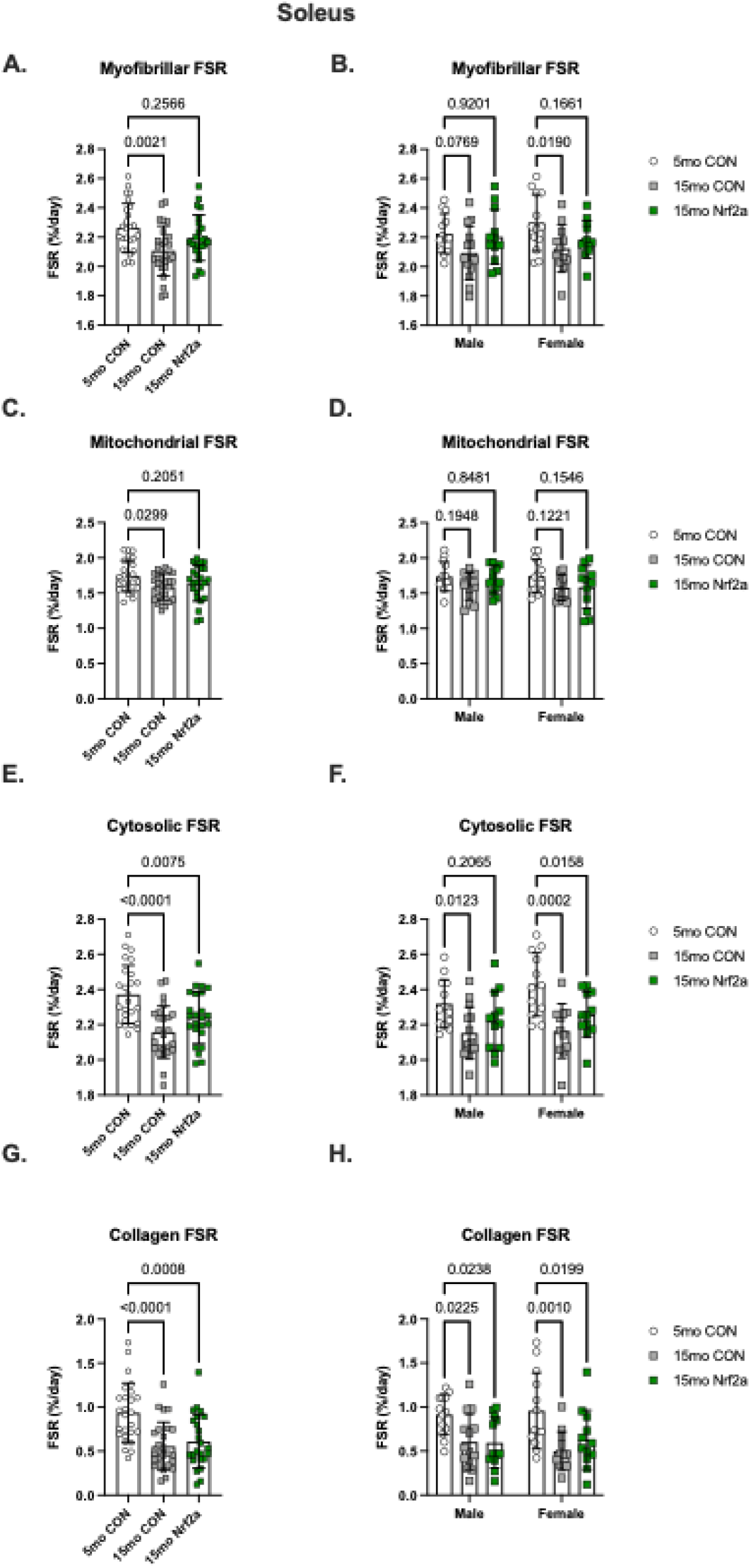
Nrf2a treatment attenuates age-related declines in FSR of soleus subfractions. 15 mo CON guinea pigs had lower FSR in each subfraction of the soleus (p=0.0021, p=0.030, p<0.0001, p<0.0001 for the myofibrillar **(A)**, mitochondrial **(C)**, cytosolic **(E)**, and collagen **(G)** subfractions, respectively). Nrf2a attenuated the decline in the myofibrillar **(A)** and mitochondrial **(C)** subfractions, but not in the cytosolic **(E)** or collagen subfractions **(G)**. Nrf2a attenuated the decline in myofibrillar FSR in both males (p=0.920) and females (p=0.166) **(B)** and attenuated the decline in cytosolic FSR in males only (p=0.207) **(F)**.

**Figure 8.**
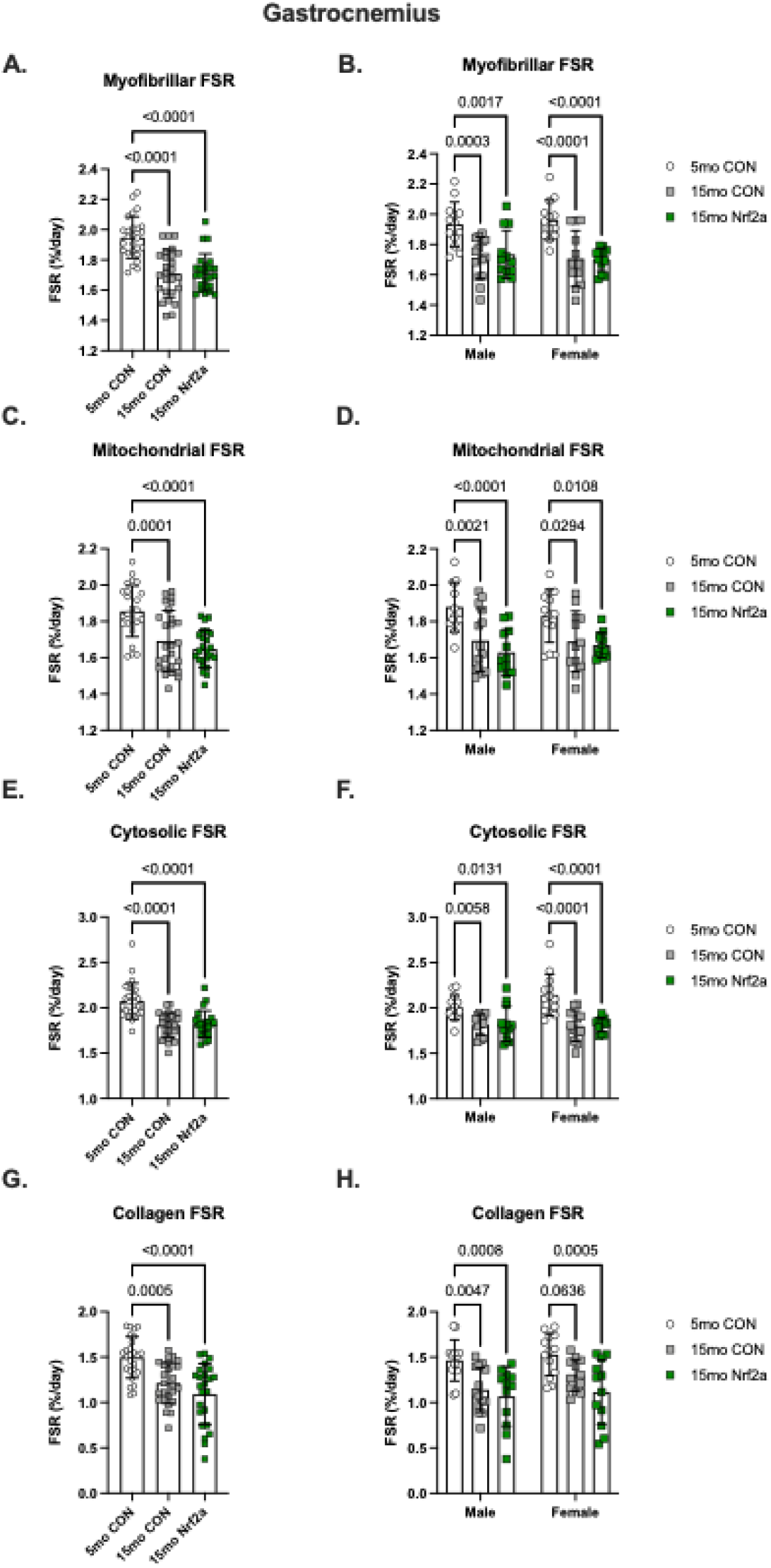
Nrf2a treatment does not attenuate the age-related decline in FSR in the gastrocnemius. 15 mo CON guinea pigs had significantly lower FSR compared to 5 mo CON guinea pigs in each subfraction (p<0.0001 for all subfractions) **(A, C, E, G)**. The FSR of all subfractions of the gastrocnemius in 15 mo Nrf2a guinea pigs were also significantly lower compared to 5 mo CON guinea pigs (p<0.0001 for all subfractions) (**A, C, E, G)**. This pattern was observed in both male and female guinea pigs **(B, D, F, H)**.

### Nrf2a does not affect protein synthesis related to proteostasis

Because protein synthesis is an essential process for both growth and proteostasis, it was necessary to discern the relative amount of protein synthesis allocated towards protein turnover (i.e. proteostasis). To do this, proteins synthesis rates were evaluated relative to the rates of cell proliferation. An increased protein synthesis rate to DNA synthesis rate ratio (PRO:DNA) suggests a greater allocation of newly synthesized proteins associated with maintaining the cellular proteome, with less dedicated to new cell proliferation. In 5 mo guinea pigs, there was no effect of Nrf2a on the PRO:DNA in the gastrocnemius or soleus (Supplemental Figure 8). Similarly, there was no difference in PRO:DNA in 15 mo guinea pigs (Figure 9). Given the constrained sample size due to loss of sample, further investigation is warranted. Given the lack of effect of Nrf2a on PRO:DNA, which is reflective of the proportion of proteins synthesized allocated to proteome maintenance, it is unsurprising that there were no differences in protein carbonyl content, a marker of protein damage, in the soleus or gastrocnemius (Supplemental Figure 9).

**Figure 9.**
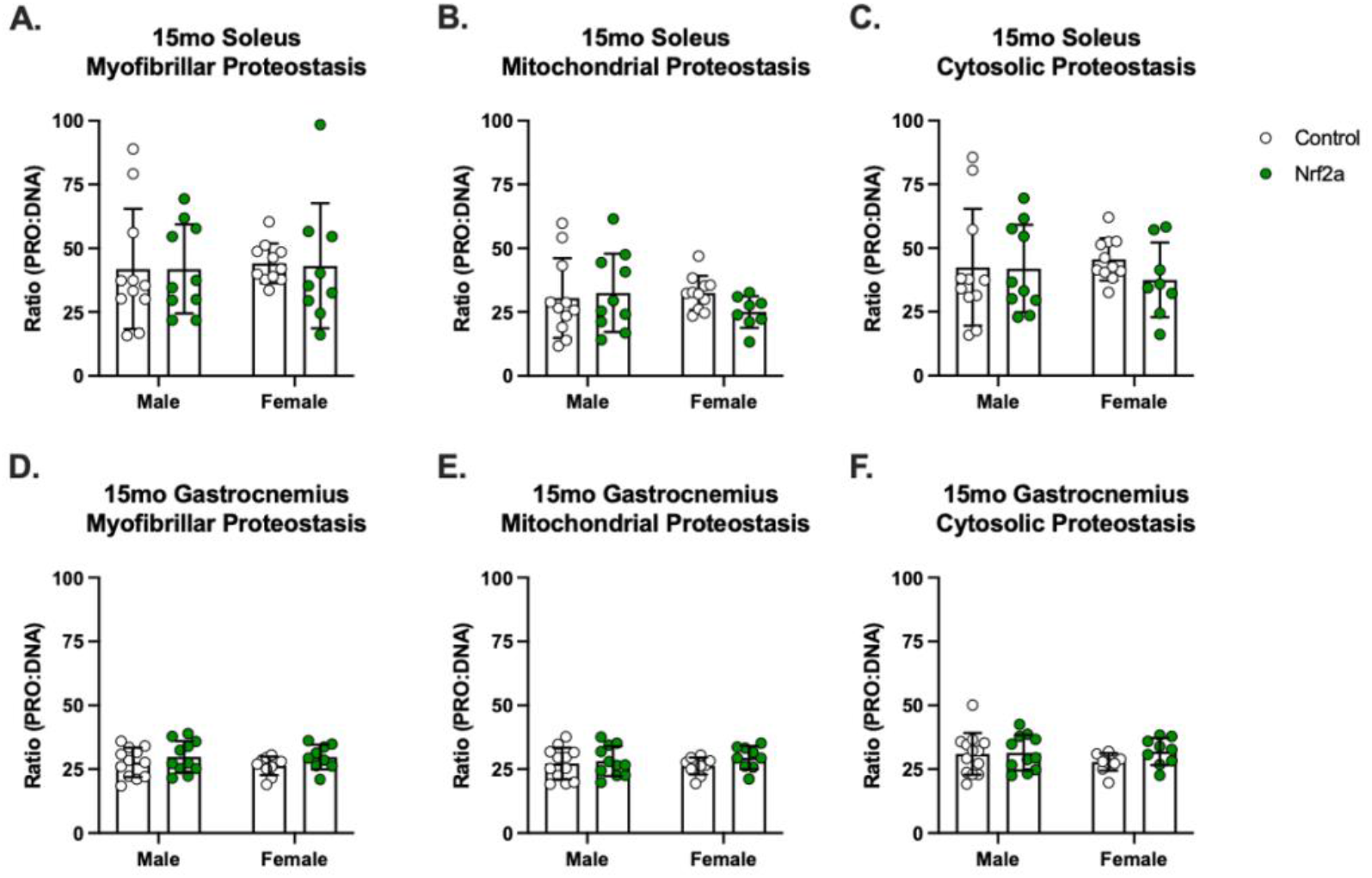
The effect of Nrf2a treatment on PRO:DNA synthesis ratios in the soleus and gastrocnemius. There was no effect of Nrf2a treatment in the myofibrillar, mitochondrial, or cytosolic subfractions on the ratio of protein to DNA synthesis in either the soleus or gastrocnemius of 15 mo male or female guinea pigs.

### The effect of Nrf2a on mobility

To determine whether or not improvements in skeletal muscle mitochondrial function and proteostasis translated to improvements in mobility, we assessed voluntary activity in a dark, enclosed area using overhead monitoring. Kaplan-Meier curves depicting the probability of sustained voluntary mobility throughout the 40-week study period. There was no statistically significant effect of Nrf2a on maintained mobility in either male or female guinea pigs. CON guinea pigs lost mobility more rapidly than Nrf2a guinea pigs (Figures 10A – B; grouped sex hazard ratio=0.713, 95% CI=0.3501 to 1.453; median ratio=1.5, CI=0.756 to 2.976; p=0.231). Further, Nrf2a males tend to have a relative increase in mobility compared to controls until about 32 weeks into the study. However, 50% of Nrf2a males lost mobility by 36 weeks, while 50% of CON males maintained mobility the entire 40-week study duration (remaining animals were censored at this time) (Figure 10A). For the majority of the study, Nrf2a females maintained their mobility compared to CON females. Approximately 50% of control females loss mobility around 16 weeks, while Nrf2a treated females sustained voluntary mobility until about 28 weeks (Figure 10B).

**Figure 10.**
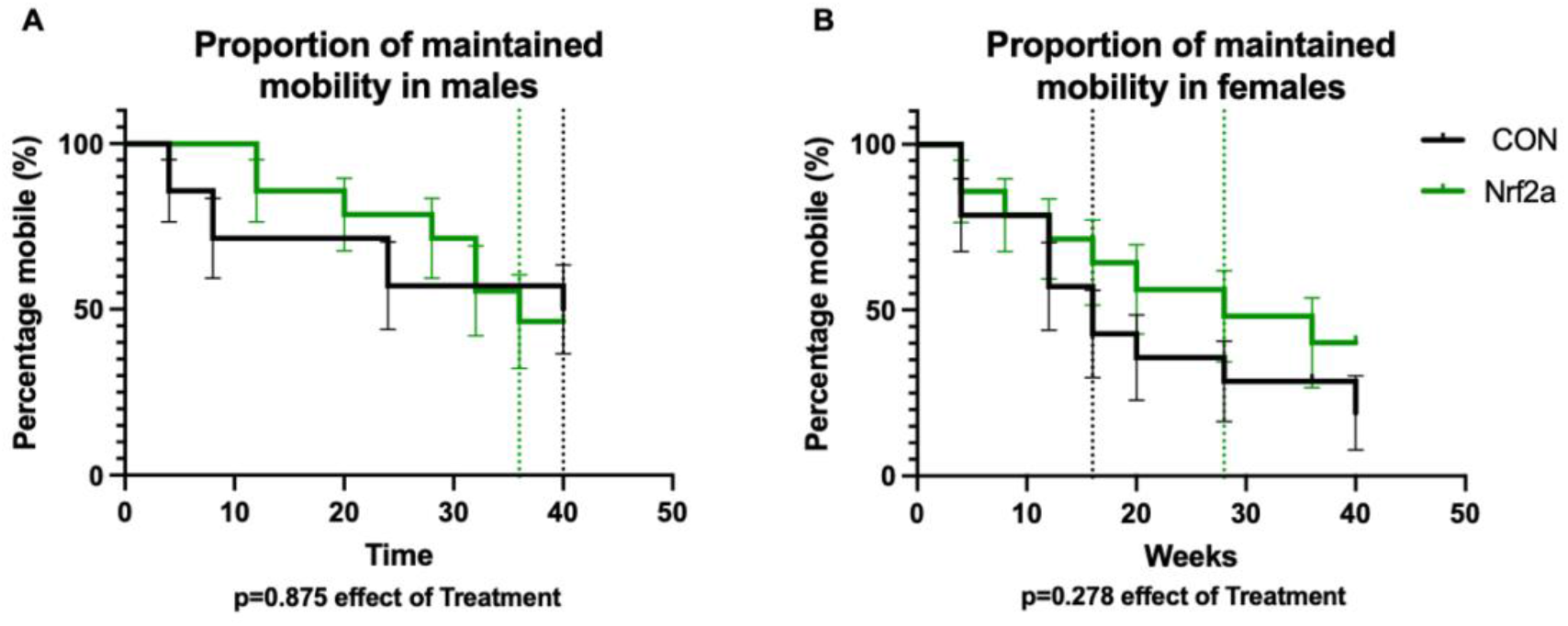
The probability of maintained mobility. There was a greater proportion of Nrf2a treated male **(A)** and female **(B)** guinea pigs that maintained mobility over the course of the study period. However, there was no statistically significant effect of Nrf2a on the probability of maintaining mobility throughout the course of the study.

## Discussion

In this study, we tested the effects of a novel phytochemical Nrf2 activator, PB125 on two hallmarks of aging implicated in musculoskeletal decline in humans: mitochondrial dysfunction and loss of proteostasis in locomotor muscle. We observed that Nrf2 activator treatment (Nrf2a) ameliorated declines in skeletal muscle mitochondrial function and protein synthesis in both male and female Hartley guinea pigs as these guinea pigs age and develop knee OA. The improvements and maintenance of mitochondrial respiration and proteostatic mechanisms may also be associated with prolonged maintenance of voluntary activity in females. Collectively, this study demonstrates the potential utility of Nrf2 activators in targeting musculoskeletal decline.

### Sex- and age/disease-related differences in mitochondrial respiration

This is the first study to measure skeletal muscle mitochondrial function in either male or female Hartley guinea pigs using high resolution respirometry. Accordingly, we first sought to characterize differences between male and female guinea pigs at 5 and 15 months of age to determine sex differences and age- and disease-related (i.e. worsening knee osteoarthritis) changes in skeletal muscle mitochondrial respiration. We found a clear sex difference in coupled and uncoupled respiration (females and lower rates of oxygen consumption than males), accompanied by decreased fatty acid supported respiration and ADP kinetics. Interestingly, these differences do not seem to be a consequence of differences in mitochondrial density and may instead reflect intrinsic differences in mitochondrial function. One study in both young and old men and women determined that there was no difference in phosphocreatine recovery post-exercise, a metric of mitochondrial capacity (Kent-Braun & Ng, 2000). However, measuring ATP production using bioluminescence revealed that mitochondria of men have greater capacity to produce ATP than that of women (Karakelides *et al*., 2010). Employing high resolution respirometry has revealed equivocal results; thus, it remains unclear whether or not females have greater oxidative capacity than males (Cardinale *et al*., 2018; Miotto *et al*., 2018). Regardless, it is essential to continue interrogating potential sex differences in mitochondrial function and changes that occur with both age and disease in both sexes.

Mitochondrial function declines with age and contributes to the aging process in humans (Short *et al*., 2005; Gonzalez-Freire *et al*., 2015; Distefano *et al*., 2017; Gonzalez-Freire *et al*., 2018). We demonstrated that both male and female Hartley guinea pigs similarly experience a decline in mitochondrial respiration as humans do. However, given the relatively early age of these guinea pigs (15 months; ∼10% of recorded maximal lifespan (Gorbunova *et al*., 2008), and ∼25% of average companion guinea pig lifespan (Quesenberry *et al*., 2021)), is difficult to ascertain if these changes are a consequence of either age, a consequence of the underlying factors that drive osteoarthritis and musculoskeletal dysfunction, or a combination of both. Other laboratory and companion animal guinea pigs do not exhibit such phenotypes as early in their lifespans (Santangelo *et al*., 2011; Musci *et al*., 2020). Notably, osteoarthritis is associated with impaired mitochondrial function and redox metabolism in degenerating joints (Loeser, 2010; Collins *et al*., 2016; Farnaghi *et al*., 2017; Collins *et al*., 2018). In the current study, both coupled and uncoupled respiration, as well as mitochondrial efficiency, declined with age/disease progression in both male and female guinea pig skeletal muscle (Figures 2A – D). There was also a decline in fatty acid supported oxidation (Figure 3C). In contrast, ADP Vmax unexpectedly increased with age in both male and female guinea pigs. Given the non-uniform changes in mitochondrial complex protein content (Supplemental Figure 7F – J), it is unclear if differences in mitochondrial density explain the age/disease-related declines in respiration. However, these data clearly demonstrate that impaired mitochondrial respiration is a characteristic of this pre-clinical model of musculoskeletal decline.

### Nrf2 activator treatment ameliorates age-related declines in mitochondrial respiration

Nrf2a treated guinea pigs had augmented mitochondrial function in 5 mo females and 15 mo males as characterized greater ADP Vmax and electron transport system capacity ETS_[CI-CIV]_. Importantly, Nrf2a attenuated age/disease-related dysfunction of Complex I and II supported coupled and uncoupled respiration and fatty acid oxidation in both sexes. Notably, Nrf2a selectively attenuated the age/disease-related decline in coupled respiration in males and uncoupled respiration in females. Nrf2a attenuated the age/disease-related declines in mitochondrial efficiency/coupling in males only. In humans, mitochondrial coupling decreases with age (Kumaran *et al*., 2005). Exercise-induced attenuation in loss of mitochondrial efficiency/coupling with age (Conley *et al*., 2013) has led researchers to speculate that improving mitochondrial efficiency may help attenuate sarcopenia (Harper *et al*., 2021). Interestingly, in the presence of rotenone, a Complex I inhibitor, there was no effect of Nrf2a, which suggests that Nrf2a improves mitochondrial respiration through improvements in Complex I function. This is consistent with data from another study that used a different Nrf2 activator, sulforaphane, and demonstrated improvements in Complex I function (Bose *et al*., 2020). The pathways underlying the effect of Nrf2 activation on mitochondrial function are not entirely understood. However, several studies have demonstrated that Nrf2 is a central mediator for improvements in mitochondrial function. Nrf2 at least partially mediates exercise-induced mitochondrial biogenesis and improvement in mitochondrial function (Merry & Ristow, 2016; D’Souza *et al*., 2020; Islam *et al*., 2020). Interestingly, both Nrf2-related redox signaling (Safdar *et al*., 2010) and Complex I function decrease with age in skeletal muscle (Kruse *et al*., 2016). Thus, Nrf2a may target a critical mechanism that contributes to age-related mitochondrial dysfunction, though the specific mechanisms by which Nrf2 activation might contribute to Complex I function remain to be elucidated.

As a master regulator of cytoprotective gene transcription, Nrf2 is a critical component of redox homeostasis. Skeletal muscle mitochondria of aged Nrf2 knock-out mice emit significantly more ROS than aged wildtype counterparts reflecting the role of Nrf2 in regulating redox balance (Kitaoka *et al*., 2019). *In vitro*, Nrf2 knock out models have compromised Complex I activity due to impairments in NADH availability (Kovac *et al*., 2015). Importantly, pyruvate dehydrogenase is a redox sensitive enzyme responsible for supplying NADH to Complex I (Fisher-Wellman *et al*., 2015). Thus, age-related increases in oxidative stress may constrain the supply of NADH to Complex I, which would explain age-associated decline in Complex I capacity and how NAD+ supplementation restores mitochondrial respiratory capacity (Kruse *et al*., 2016; McElroy *et al*., 2020). In our study, Nrf2a increased mitochondrial function, particularly in Complex I, which may have been mediated by improved cellular redox regulation. However, future studies will need to more rigorously investigate the effect of Nrf2a on redox homeostasis.

Another potential mechanism by which Nrf2a enhanced mitochondrial function is through greater mitochondrial protein turnover. There was no consistent age- or treatment-related effect on mitochondrial protein content in the soleus or gastrocnemius (Supplemental Figure 7). However, there was an age/disease-related decline in mitochondrial biogenesis, suggesting that, in order to maintain mitochondrial density, degradation of mitochondrial proteins (i.e. mitophagy or ubiquitin dependent degradation of mitochondrial proteins) also declined. Impaired mitophagy contributes to mitochondrial dysfunction and disease in humans (Ryu *et al*., 2016; Gouspillou *et al*., 2018; Newman & Shadel, 2018). Importantly, Nrf2a attenuated the age/disease-related decline in mitochondrial protein synthesis, suggesting that declines in degradation/mitophagy may have also been attenuated, though we did not directly measure this. As such we posit that mitochondrial protein turnover, which is essential for maintenance of overall mitochondrial function (Szczepanowska & Trifunovic, 2021), was maintained in 15 mo Nrf2a guinea pigs compared to 15 mo CON guinea pigs in this study. Others have also demonstrated that Nrf2 activators play a role in modulating mitochondrial protein turnover. In *C. elegans* the Nrf2 homolog mediated Tomatidine-induced (a Nrf2 activator) mitophagy (Fang *et al*., 2017). Our group has demonstrated that Protandim, also a phytochemical Nrf2 activator, enhanced mitochondrial protein turnover in wheel running rats (Bruns *et al*., 2018). Thus, Nrf2 activation seems to preserve mitochondrial protein turnover in 15 mo guinea pigs while turnover may have declined in 15 mo CON guinea pigs.

### Nrf2a attenuates components of protein homeostasis

Decline in mechanisms to maintain proteostasis (which includes not only protein synthesis and degradation, but also chaperone-mediated folding and protein trafficking (Noack *et al*., 2014)) contributes to age-related musculoskeletal dysfunction (Kaushik & Cuervo, 2015; Santra *et al*., 2019). There is limited insight on the effect of age on protein homeostasis in humans, though basal protein synthesis appears to be unchanged with age in humans (Volpi *et al*., 2001; Brook *et al*., 2016). Moreover, differences between men and women with regard to the decline in skeletal muscle proteostasis remains unclear. While men generally have greater muscle mass than women, men also lose muscle mass faster and muscle strength to a greater degree; however, women are less fatigue resistant (thoroughly reviewed in (Gheller *et al*., 2016). In the present study, we documented the age-related decline in protein synthesis in all subfractions of the soleus and gastrocnemius muscles of both male, which we observed in our previous study (Musci *et al*., 2020), and female Hartley guinea pigs. There were no sex differences in fractional synthesis rates in either muscle. This is the first study to characterize age-related declines in protein synthesis in female guinea pigs. It is important to note, however, that while we documented age-related differences in long-term protein synthesis rates to minimize the bias of faster turning over proteins (Miller *et al*., 2015), it is possible that our approach may still not accurately determine differences in fractional synthesis rates between ages if the protein pools subject to turnover (i.e. the dynamic protein pools) are not the same between the 5 mo and 15 mo guinea pigs. As recently demonstrated by Abbott and Lawrence and colleagues, the dynamic protein pool declines with age and thus obscures the fractional synthesis rates and biases towards aged animals having lower synthesis rates (Abbott *et al*., 2021). The approach the authors employed is both novel and unique, but raises important considerations when evaluating the effect of age or interventions on protein turnover in the future. Employing such an approach may also help reconcile differences in observations on the effect of age on protein turnover between species (Volpi *et al*., 2001; Miller *et al*., 2019; Musci *et al*., 2020) and more accurately describe the age-related effects on protein kinetics. Importantly, we agree with the authors that adopting such a rigorous approach in the future will provide better guidance as to how to improve proteome integrity and maintain the dynamic protein pool with age.

In the present study, Nrf2a attenuated the age/disease-related declines in myofibrillar and mitochondrial protein synthesis rates in the soleus in both males and females. Interestingly, Nrf2a had no attenuating effect on the age/disease-related declines of protein synthesis in any subfraction of the gastrocnemius. One driving factor of protein synthesis is cellular proliferation (Eden *et al*., 2011). Thus, to discern protein synthesis dedicated to proliferation as opposed to proteome maintenance, we made simultaneous measurements of DNA synthesis rates to provide insight about the proportion of protein synthesis dedicated towards newly synthesized proteins compared to proteome maintenance (Miller *et al*., 2014). There was no difference in the allocation of protein synthesis to proteome maintenance in the soleus or gastrocnemius in either male or female guinea pigs. Moreover, Nrf2a had no effect on protein carbonylation levels in either the soleus or gastrocnemius. These data are in contrast with our previous studies demonstrating that other Nrf2 activators promote proteome maintenance *in vitro* and *in vivo* in both rats (Bruns *et al*., 2018) and humans (Konopka *et al*., 2017). Importantly, interventions that activate mechanisms maintaining proteostasis are linked to healthspan extension in a variety of organisms (Pride *et al*., 2015; Hamilton & Miller, 2017; Sands *et al*., 2017). Thus, while Nrf2a attenuated the decline in protein synthesis in the present study, Nrf2a did not increase the proportion of proteins synthesized for proteome maintenance.

The mechanisms by which Nrf2a attenuated the decline in protein synthesis are not entirely clear. However, alleviating energetic constraints through enhanced mitochondrial function is a likely candidate to explain some of these improvements. Protein turnover is energetically demanding, accounting for nearly 35% of basal metabolism (Waterlow, 1984; Rolfe & Brown, 1997; Bier, 1999). Age-related impairments in mitochondrial function consequentially constrain the amount of energy dedicated to proteostasis. Mitochondrial dysfunction precedes the loss of proteostasis in skeletal muscle, which leads to declines in function (Ben-Zvi *et al*., 2009; Gaffney *et al*., 2018). Moreover, other interventions that attenuate the decline in or improve mitochondrial function, also improve proteostatic mechanisms and preserve overall muscle function. For example, maintaining physical activity and caloric restriction in rodents delays declines in mitochondrial function as well as skeletal muscle function (Zangarelli *et al*., 2006; Stolle *et al*., 2018), a similar observation made in masters athletes (Zampieri *et al*., 2015). While both exercise and caloric restriction have broad effects, more targeted interventions focused on improving mitochondrial function also report a similar phenomenon: enhancing mitochondrial function delays skeletal muscle dysfunction (Gaffney *et al*., 2018; Campbell *et al*., 2019). This observation occurs in other tissues as well. Increasing mitochondrial proteostasis decreases proteotoxic amyloid aggregation in cells, increasing fitness and lifespan in *C. elegans* (Sorrentino et al., 2017). These studies emphasize the importance of mitochondrial respiration and the production of ATP to facilitate proteostatic mechanisms. In humans, aerobic exercise improves mitochondrial function through mitochondrial remodeling and improves skeletal muscle function (Greggio *et al*., 2017). Altogether, our data support the posit that Nrf2a-mediated improvements in mitochondrial respiration alleviated constraints in energy which led to greater amount of ATP available to support proteostasis.

Another mechanism by which Nrf2a may have attenuated declines in skeletal muscle proteostasis is through the mitigation of inflammation and oxidative stress, which can have deleterious effects on protein turnover, particularly protein synthesis. Protein synthesis, at rest, appears to be no different between young and old individuals (Volpi *et al*., 2001; Brook *et al*., 2016). However, age-related inflammation and oxidative stress can blunt the anabolic response to stimuli such as exercise or feeding. This concept, termed anabolic resistance, is a contributor to age-related musculoskeletal dysfunction and appears to blunt the anabolic response to resistance exercise training (Cuthbertson *et al*., 2005; Wilkes *et al*., 2009; Burd *et al*., 2012; Brook *et al*., 2016). Interventions designed to mitigate age-related increases oxidative stress or inflammation seem to improve skeletal muscle anabolic responses to exercise (Trappe *et al*., 2002), feeding (Rieu *et al*., 2009; Smiles *et al*., 2019), and insulin (Rivas *et al*., 2016). Importantly, Nrf2a stimulates transcription of endogenous antioxidant and anti-inflammatory genes (Hybertson *et al*., 2011; Hybertson *et al*., 2019). Thus, it is possible that Nrf2a treatment ameliorated oxidative stress and inflammation and improved the anabolic response to feeding. However, because we measured cumulative protein synthesis over 30 days, rather than acutely in response to an anabolic stimulus such as feeding, we cannot determine if there were any changes specifically in the anabolic response to feeding. Future studies should investigate the efficacy of this particularly Nrf2a, PB125, on abrogating inflammation and oxidative stress and, in acute settings, determine whether Nrf2a enhances the anabolic response to feeding or activity. In the present study, there was no observed effect of treatment on ROS emission or protein oxidation. However, our lab has previously demonstrated that another Nrf2 activator increases antioxidant protein expression (Reuland *et al*., 2013) and augmented protein synthesis related to proteostasis in skeletal muscle of rats in response to wheel running exercise (Bruns *et al*., 2018). Thus, treatment with Nrf2 activators may represent a class of interventions that augment adaptation to acute stressors (Musci *et al*., 2019).

### Future Directions

The improvements in mitochondrial respiration and proteostasis did not translate to sustained improvements in mobility. However, it is important to note that our measure of mobility is only one metric of musculoskeletal function. Additionally, musculoskeletal function is not the only factor that dictates mobility. Thus, it is still important to assess other and more specific components of musculoskeletal function, as mitochondrial function is a strong determinant in physical function such as gait speed and grip strength in humans (Gonzalez-Freire *et al*., 2018).

Another observation that warrants further investigation are the sex-specific effects of Nrf2a. Other interventions involving Nrf2 activators have also demonstrated sex-specific effects. The Interventions Testing Program reported that treatment with the Nrf2 activator Protandim extended median lifespan in heterogenous male mice, but not females (Strong *et al*., 2016). Our lab has also previously demonstrated that Protandim only improved myofibrillar proteostasis in men (Konopka *et al*., 2017). Other healthspan promoting interventions, such as a metformin and rapamycin, also have sex specific effects (Strong *et al*., 2016). We have begun interrogating these sex specific effects through the use of kinetic proteomics (Wolff *et al*., 2019; Wolff *et al*., 2021). Moving forward, it will be necessary to interrogate these sex differences and the implications they have on the efficacy of Nrf2a to attenuate musculoskeletal decline. Females experience sarcopenia at a similar prevalence as males worldwide (Shafiee *et al*., 2017), future investigation of targeting Nrf2 for musculoskeletal aging should include investigation of the mechanisms likely to underlie our sex-specific responses including interaction of reproductive hormones with Nrf2 signaling.

### Conclusions

Musculoskeletal dysfunction is a primary contributor to disability and dependence with age. There are few existing interventions that effectively mitigate the decline in skeletal muscle function with age. In this study, we further characterized a model for musculoskeletal dysfunction measuring mitochondrial function and protein synthesis in both male and female Hartley guinea pigs. Moreover, we tested a potential healthspan-extending phytochemical compound PB125, which is currently in the NIA-ITP (https://www.nia.nih.gov/research/dab/interventions-testing-program-itp/compounds-testing), on mitochondrial function and proteostasis in this pre-clinical guinea pig model of musculoskeletal decline. We found that this compound improved mitochondrial function and attenuated declines in protein synthesis, mechanisms that likely mediate improvements in function and longevity. This project adds to the growing literature that supports the use of Nrf2 activators to improve organismal health. The data from this study provide mechanistic insight by which a readily translatable intervention could mitigate age-related musculoskeletal decline.

## Acknowledgments

We would like to thank Dr. Ann Hess of the Statistics Department at Colorado State University for overseeing and guiding the statistical analysis of this project. Pathways Bioscience provided the PB125 compound for this project. This work was funded by NIH grant R21 AG054713-02 awarded to KLH and KSS and supported by the ACSM NASA Space Physiology Grant awarded to RVM.

## Author Contributions

Study design: RVM, KMA, BFM, MAJ, JMM, BMH, KSS, KLH. Data collection: RVM, KMA, MAW, ZV, MFA, SB, MC, TJ, TEK, RM, TN, JS, SW, MDM, QZ. Data analysis: RVM, KMA, KSS, KLH. Manuscript preparation: RVM, KLH. All authors approved of the final manuscript.

