## Supplemental Data for "Phytochemical Nrf2 activator attenuates skeletal muscle mitochondrial dysfunction and impaired proteostasis in a preclinical model of musculoskeletal aging"

**Supplemental Table 1** Summary of high resolution respirometry experiment using Ouroboros Oxygraph-2000. A table highlighting both SUIT 1 and SUIT 2 protocols used in the study. Further explanations of these respiratory states and measurements can be found in Pesta & Gnaiger, 2011 or at the Ouroboros website: www.bioblast.at.

Abbreviations: P – pyruvate; G – glutamate; M – malate; S – succinate; Oct – octanoylcarnitine; ADP – adenosine diphosphate; ATP – adenosine triphosphate; Rot – rotenone; FCCP – Trifluoromethoxy carbonylcyanide phenylhydrazone; Ama – Antimycin A; ETS – electron transport system; Sub – priorly added substrates (i.e. pyruvate, glutamate, malate, octanoylcarnitine, and succinate); CI – Complex I; CII – Complex II; CIII – Complex III; CIV – Complex IV; CV – Complex V or ATP Synthase; ROX – residual oxygen consumption; RCR – respiratory control ratio; NADH – reduced nicotinamide adenine dinucleotide;

**Supplemental Figure 1** The concentration and stability of phytochemical components and Nrf2 content in guinea pig skeletal muscle. The concentration of luteolin **(A)**, carnosol **(B)**, and withaferin A **(C)** in plasma after dosing guinea pigs with 8, 24, or 40 mg/kg of PB125 from 0 to 120 min. The stability of the phytochemical components suspended in vehicle at room temperature and in 4 $^{o}$C **(D)**. Nrf2 content was significantly increased ($p=0.009$) as a result of Nrf2a treatment in the gastrocnemius of guinea pigs **(E)**.

**Supplemental Figure 2** Nrf2a treatment does not affect growth of guinea pigs. Growth charts for male **(A)** and female **(B)** guinea pigs. There was a significant effect of Age ($p<0.0001$) and Sex ($p<0.0001$) on guinea pig body mass **(C)**. There was a significant decrease ($p<0.0001$) in skeletal muscle DNA synthesis with age in the soleus **(D)** and gastrocnemius **(E)**. Female guinea pigs also had a higher ($p=0.005$) rate of DNA synthesis in the gastrocnemius **(E)**. There was also a significant interaction between Age and Sex ($p=0.0062$) **(E)**.

**Supplemental Figure 3** Age-related changes in skeletal muscle mass. There was a significant effect of Age ($p<0.0001$) in soleus, gastrocnemius, plantaris, and tibialis anterior muscles **(A – D)**. Male guinea pigs had greater ($p<0.0001$) skeletal muscle mass compared to females **(A – D)**. There was no effect of Age in either the relative (mg of muscle/g of body mass) soleus **(E)** or plantaris **(G)** muscles, however there was a significant ($p=0.005$) negative effect of Age on the gastrocnemius **(F)** and tibialis anterior **(H)**.

**Supplemental Figure 4** Cytochrome C Control Factor Scatterplots**.** Scatterplots and regression line relating Cytochrome C Control Factor to coupled respiration in Suit 1 **(A)** and Suit 2 before **(B)** and after **(C)** a limit of 0.30 was implemented to establish O2K trials to exclude due to over permeabilization.

**Supplemental Figure 5** Nrf2a treatment had no effect on ROS emission in male or female guinea pigs. Nrf2a had no effect on ROS emission during LEAK (i.e. State 2 respiration) **(A)** or State 3 respiration in the presence of sub-saturating amounts (0.5 mM) of ADP **(B)**.

**Supplemental Figure 6** ADP Titration Curves. Averages of ADP Vmax and Km of male and female guinea pigs **(A-D)** and representative best-fit curves for averaged values **(E – H)**.

**Supplemental Figure 7** Nrf2a treatment had no effect on mitochondrial proteins. There were no Age-, Sex-, or Treatment-related effects on mitochondrial protein content in Complexes I – V in the gastrocnemius **(A – E)**. There was a significant effect of Age (p=0.0126) and Sex (p=0.0789) on Complex I protein content in the soleus **(F)**. There was also an significant effect of Age in Complex III (p=0.013) **(H)** but not in Complex IV protein content (p=0.058) **(I)**.

**Supplemental Figure 8** There was no effect of Nrf2a treatment on PRO:DNA synthesis ratios in 5 mo guinea pigs. There was no significant of Age or Treatment in any of the subfractions of the soleus **(A – D)** or gastrocnemius **(E – H)**.

**Supplemental Figure 9** The effect of Sex, Age, and Treatment on protein carbonyl concentration in skeletal muscle. There was no effect of Age, Sex, or Treatment on protein carbonyl concentration in the soleus **(A)**. There was no significant increase (p=0.0814) in protein concentration in the gastrocnemius **(B)**.

| Respiratory State | Substrates | Description |
|---|---|---|
| **SUIT 1 – A Complex I supported ADP titration evaluating ADP sensitivity as well as maximal respiration, followed with uncoupled respiration.** | | |
| **State 2[PGM]** | 5 mM pyruvate (P), 10 mM glutamate (G), 0.5, mM malate (M) | LEAK respiration in the presence of Complex I substrates. The amount of oxygen consumption as a consequence of protons leaking across the membrane. |
| **State 3[PGM]** | Previous steps + addition of ADP such that [ADP] progressively increased as follows: 0.1 mM, 0.175 mM, 0.25 mM, 1 mM, 2 mM, 4 mM, 8 mM, 12 mM, 20 mM, to 24 mM. | Respiration associated with ATP production. NADH is generated through the oxidation of PGM, donating electrons to Complex I, and creating a proton gradient where ADP is the limiting factor for ATP production. The amount of oxygen consumed is a consequence of ADP being phosphorylated to ATP. Initial titrations are sub-saturating concentrations of ADP increasing to maximal ADP stimulated respiration. |
| **State 3[PGM + S]** | Previous step + 10 mM succinate (S) | Same as previous step but now both NADH and $FADH_2$ are generated through the oxidation of Complexes I (PGM) and II (S) substrates yielding maximal ADP stimulated respiration. |
| **ETS [CI – CIV]** | Previous step + 1.0 mM FCCP | FCCP is a protonophore allowing hydrogen ions through the inner-mitochondrial membrane, uncoupling the proton gradient from ATP production through ATP synthase. This reflects the maximal or reserve capacity of Complexes I – IV, unconstrained from ATP synthase activity. |
| **ETS [CII – CIV]** | Previous step + 5 µM rotenone (Rot) | Rotenone is an inhibitor of Complex I. Therefore, uncoupled oxygen consumption is indicative of Complex II – IV capacity. |
| **ROX** | 2.5 µM Antimycin A (Ama) | Antimycin A inhibits Complex III thus shutting down the electron transfer pathway. Residual oxygen consumption (ROX) is the remaining oxygen consumed from other pathways independent of Complex IV. |
| **SUIT 2 – A protocol measuring coupled and uncoupled respiration as well as ROS emission simultaneously in the presence of fatty acids.** | | |
| **State 2[PGM + Oct]** | 5 mM pyruvate (P), 10 mM glutamate (G), 0.5, mM malate (M), 0.2 mM octanoylcarnitine (Oct) | LEAK respiration in the presence of Complex I and II substrates including both carbohydrates and fatty acids. The amount of oxygen consumption as a consequence of protons leaking across the membrane. |
| **State 2[PGM + Oct + S]** | Previous step + 10 mM succinate (S) | Same as previous step with the addition of the Complex II substrate, succinate (S). This is considered to stimulate maximal LEAK respiration as well as generation of reactive oxygen species (ROS). |
| **State 3[Sub + 0.5D]** | Previous step + 0.5 mM ADP | Respiration associated with ATP production. NADH and $FADH_2$ is generated through the oxidation of PGMS and octanoylcarnitine donating electrons to Complexes I and II, and creating a proton gradient where ADP is the limiting factor for ATP production. The amount of oxygen consumed is a consequence of ADP being phosphorylated to ATP with a sub-saturating bolus of ADP. |
| **State 3[Sub + 1.0D]** | Previous step + 0.5 mM ADP | Same as previous step. The final concentration of ADP is higher but still sub-saturating stimulated respiration. |
| **State 3[Sub + 6.0D]** | Previous step + 5 mM ADP | Same as previous step, however, the final concentration of ADP (6.0 mM) is a saturating does, maximally stimulating mitochondrial respiration and ATP production. |
| **State 3[Sub + D – CI]** | Previous step + 5 µM rotenone (Rot) | Rotenone is an inhibitor of Complex I. Oxygen consumption is due to maximally stimulated ADP respiration of Complexes II – IV. |
| **ETS[Sub + D – CI]** | Previous step + 1.0 mM FCCP | FCCP is a protonophore allowing hydrogen ions through the inner-mitochondrial membrane, uncoupling the proton gradient from ATP production through ATP synthase. This is the maximal or reserve capacity of the mitochondria for Complexes II – IV. |
| **Calculated Ratios** | **Calculation** | **Description** |
| **Respiratory Control Ratio (RCR)** | State 3[PGM] / State 2[PGM] | The ratio of maximally stimulated ATP production to LEAK respiration. Respiratory Control Ratio (RCR) is a metric of mitochondrial coupling efficiency. A higher ratio is indicative of greater mitochondrial efficiency of oxygen consumption coupled to ATP production. |

**Supplementary Table 1**

#### Supplementary Figure 1

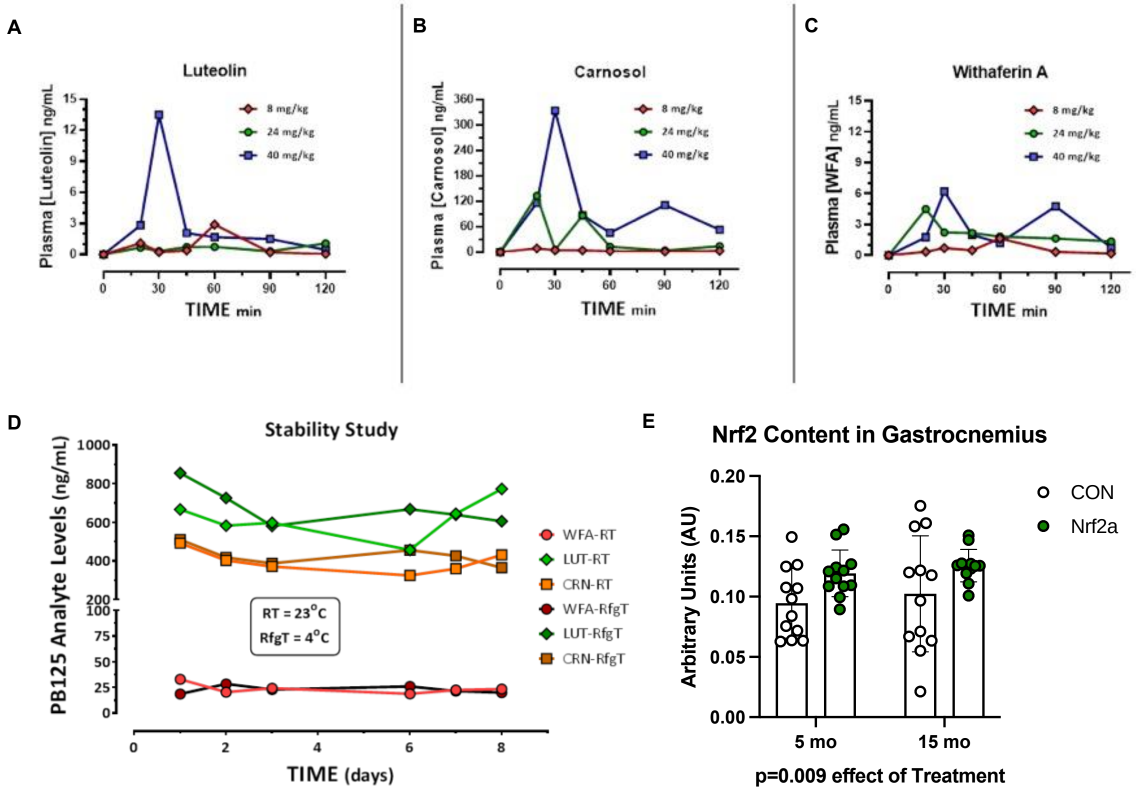

#### Supplementary Figure 2

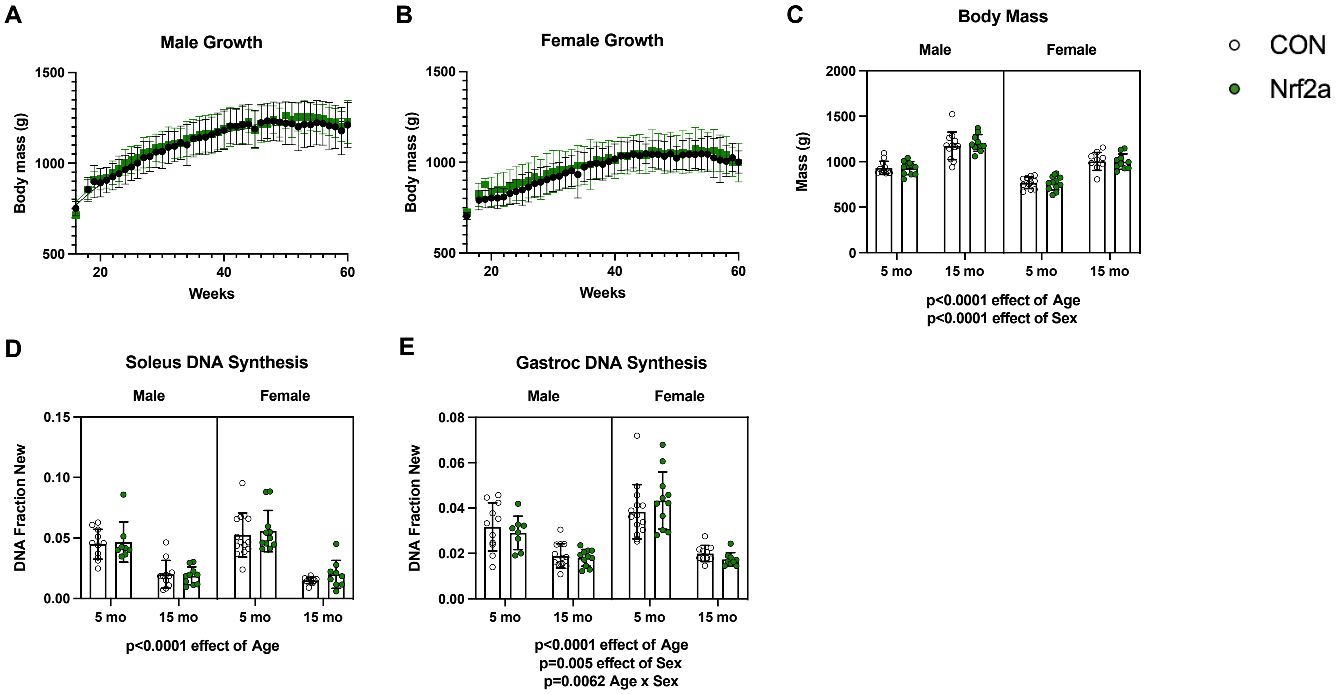

#### Supplementary Figure 3

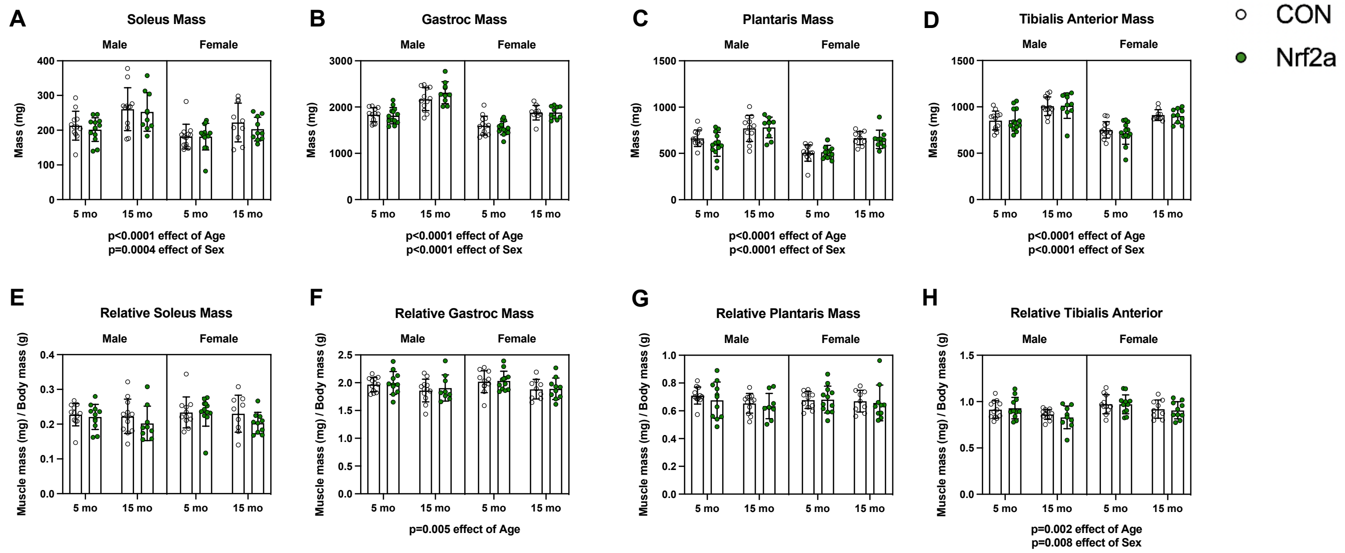

**Supplementary Figure 4**

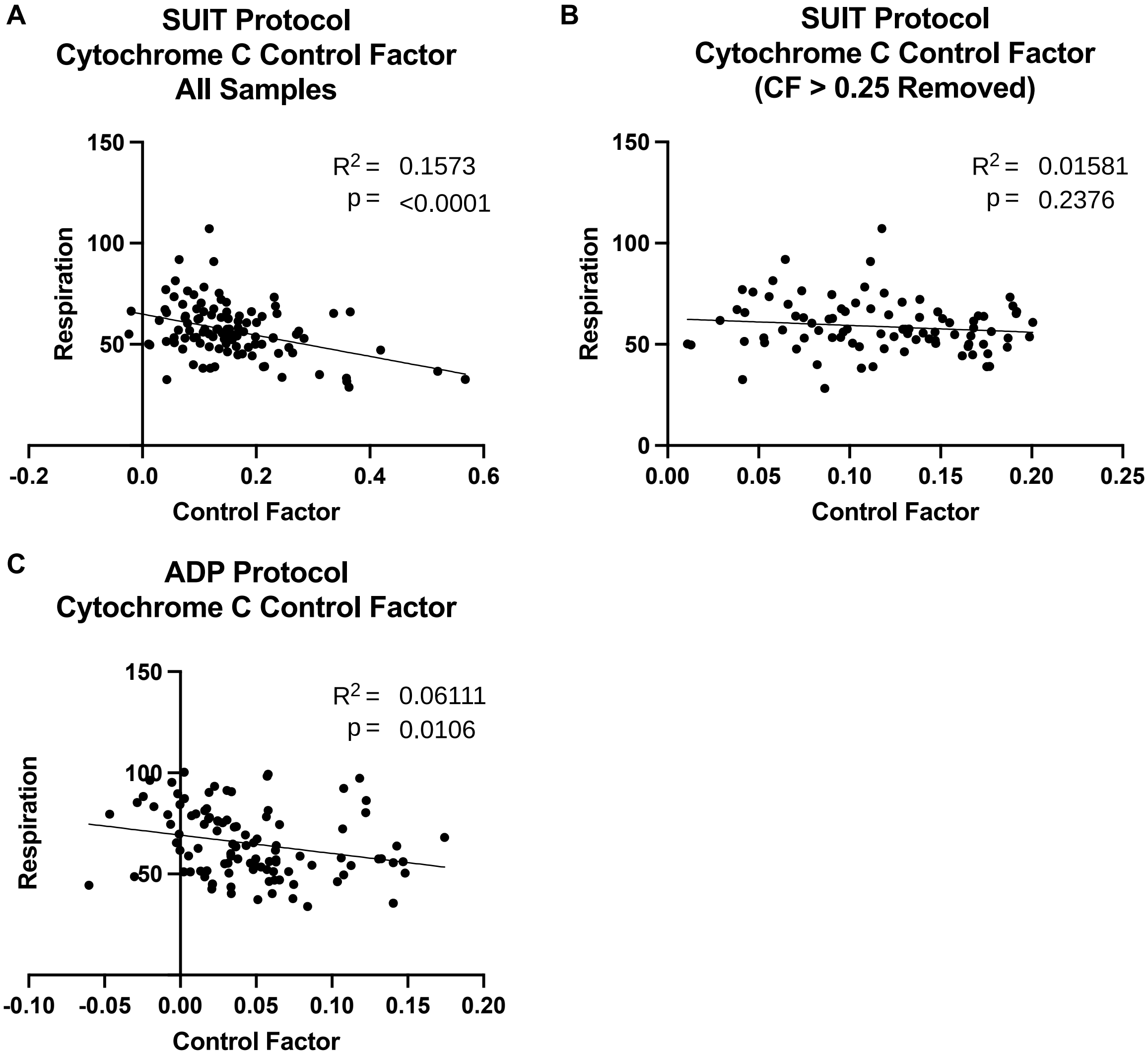

**Supplementary Figure 5**

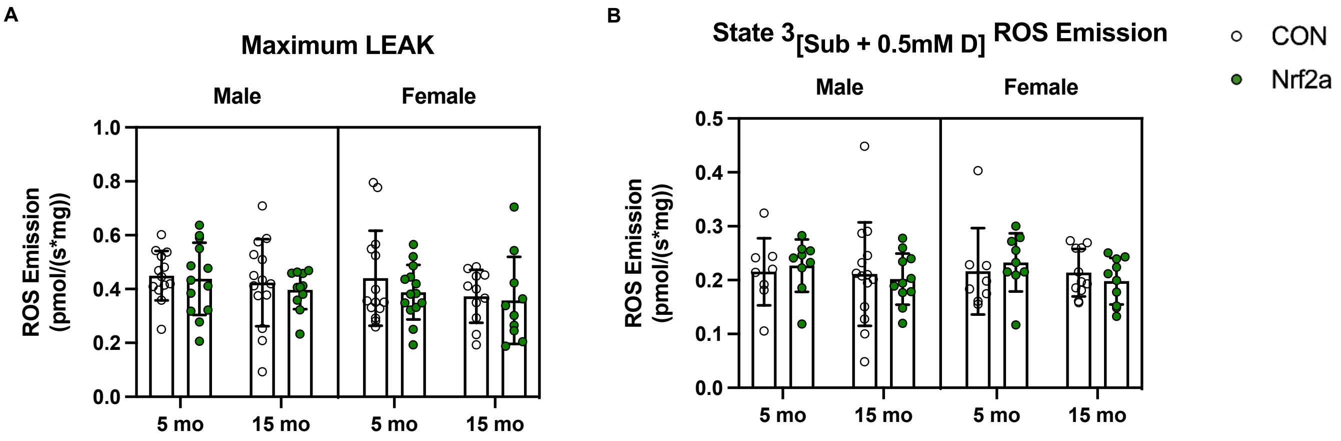

#### Supplementary Figure 6

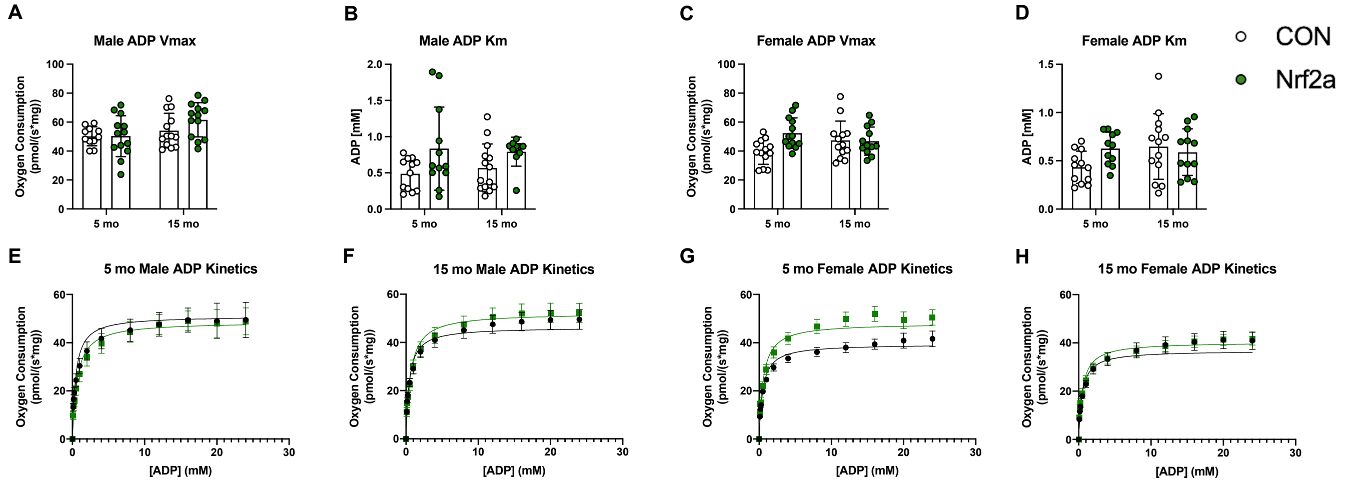

### Supplementary Figure 7

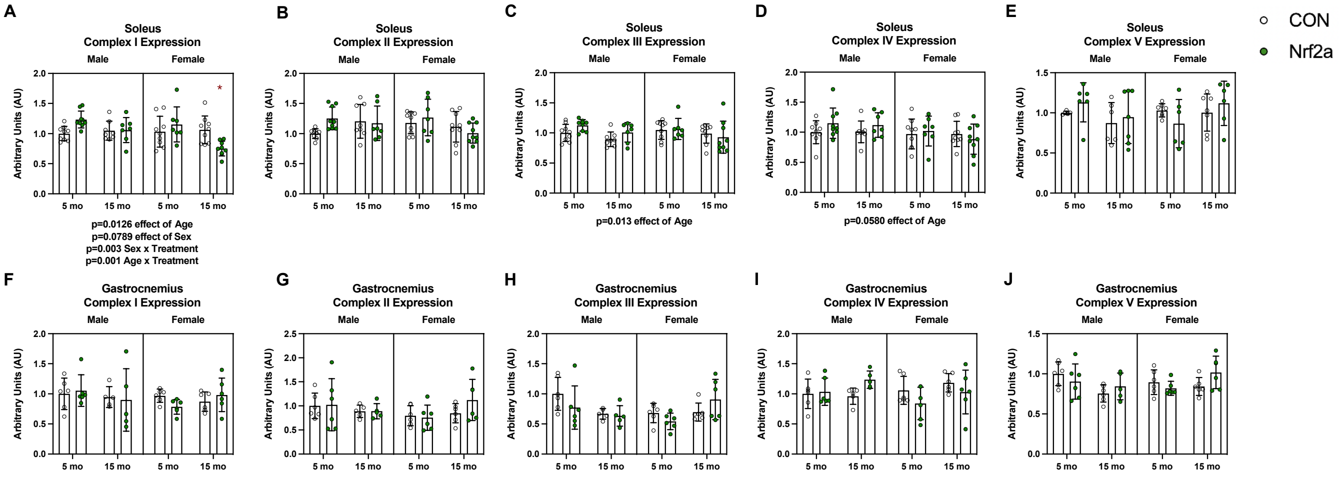

**Supplementary Figure 8**

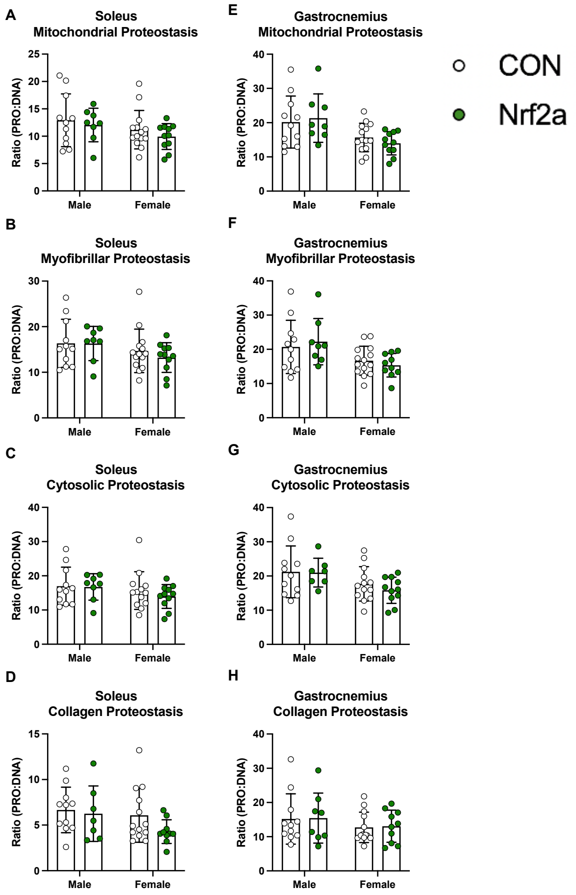

**Supplementary Figure 9**

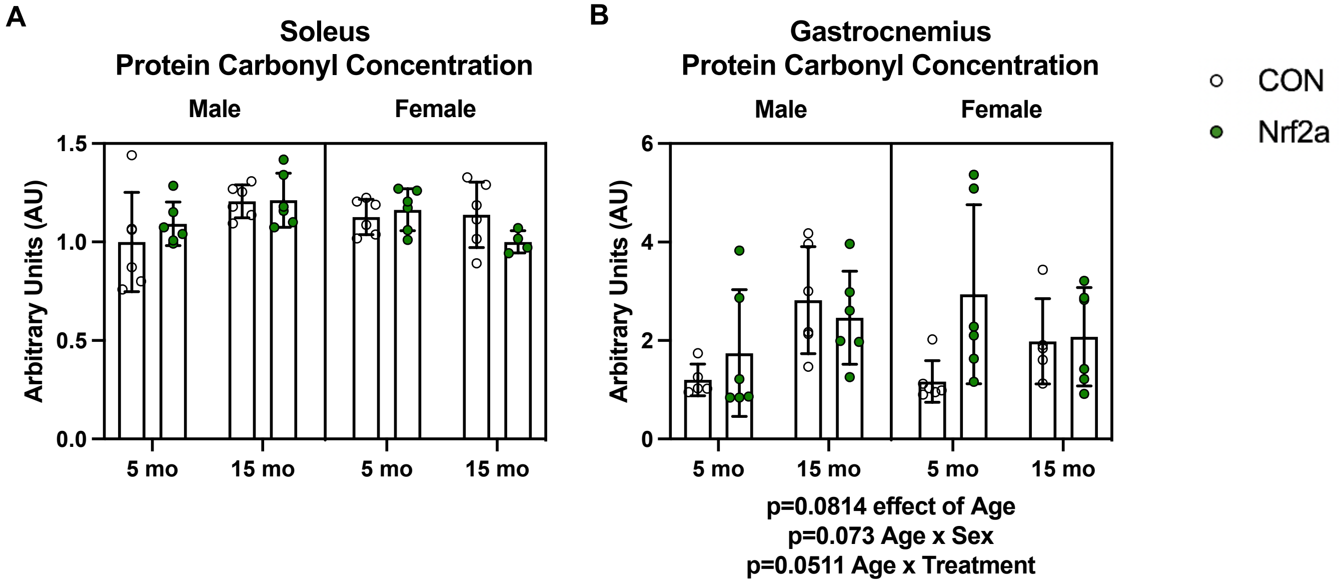
